# Distinct cellular origins and differentiation process account for distinct oncogenic and clinical behaviors of leiomyosarcomas

**DOI:** 10.1101/2020.10.23.352336

**Authors:** Elodie Darbo, Gaëlle Pérot, Lucie Darmusey, Sophie Le Guellec, Laura Leroy, Laëtitia Gaston, Nelly Desplat, Noémie Thébault, Candice Merle, Philippe Rochaix, Thibaud Valentin, Gwenaël Ferron, Christine Chevreau, Binh Bui, Eberhard Stoeckle, Dominique Ranchere-Vince, Pierre Méeus, Philippe Terrier, Sophie Piperno-Neumann, Françoise Collin, Gonzague De Pinieux, Florence Duffaud, Jean-Michel Coindre, Jean-Yves Blay, Frédéric Chibon

## Abstract

In leiomyosarcoma (LMS), a very aggressive disease, a relatively transcriptionally uniform subgroup of well differentiated tumors has been described and is associated with poor survival. The question raised how differentiation and tumor progression, two apparently antagonist processes, coexist and allow tumor malignancy. We first identified the most transcriptionally homogeneous LMS subgroup in three independent cohorts, which we named ‘hLMS’. The integration of multi-omics data and functional analysis suggests that hLMS originate from vascular smooth muscle cells and show that hLMS transcriptional program reflects both modulation of smooth muscle contraction activity controlled by MYOCD/SRF regulatory network and activation of the cell cycle activity controlled by E2F/RB1 pathway. We propose that the phenotypic plasticity of vascular smooth muscle cells coupled with MYOCD/SRF pathway amplification, essential for hLMS survival, concomitant with PTEN absence and *RB1* alteration, could explain how hLMS balance this uncommon interplay between differentiation and aggressiveness.

---

Cell differentiation is often associated with malignancy in solid tumors: in general, the more differentiated a tumor is, the less it is aggressive. Leiomyosarcoma (LMS), a rare (11% of adult soft tissue sarcomas [1]) and very aggressive (50% of patients relapse [2] with a median survival of 12 months) mesenchymal malignancy, challenges this concept.

Previous studies [3–10] characterized a subgroup of well differentiated LMS showing either equal [3,5,7], better [6,8,9] or worse [10] prognosis than the remaining LMS. These results relied on very different sample repositories (size, uterine LMS, metastatic and primary tissues), implying discrepancies between the reported subtypes. However, they demonstrated the existence of molecular subtypes that should be detected when considering treatment options. Current first-line treatment involves wide surgical resection for localized LMS or anthracycline-based chemotherapies for metastatic tumors, since neither targeted therapy [11] nor immunotherapy [12] have demonstrated any major therapeutic effects until now. However, the development of targeted therapies in LMS is challenging since their oncogenesis is still poorly understood, although genomic alterations are now well described, particularly the highly recurrent alterations of p53, RB1 and PTEN [10,13,14].

In the present study, we focus on these well differentiated LMS and aim at understanding how highly specialized cells keep or acquire their propensity to proliferate and migrate. *In fine*, we hope to detect potential therapeutic vulnerabilities, which could pave the way for a new treatment option. Moreover, previous studies highlighted the relatively homogeneous transcriptional behavior of well differentiated LMS, which we suspect arises from a unique driver event, as reported by Watson and colleagues for other sarcomas with a strong chimeric driver oncogene [15]. We thus hypothesize that a common oncogemc event might underpin this homogeneous transcriptome. This could not only shed light on the biology of these LMS but also improve patient stratification and provide a therapeutic opportunity.

## Methods

### EXPERIMENTAL MODEL AND SUBJECT DETAILS

#### Human samples

Among samples used in the training cohort, 278 out of the 387 complex genetics sarcomas [16–19], the 60 GIST [20], and the 58 synovial sarcomas [21] are part of cohorts previously described (**Table S1**). All samples used in this cohort were part of the CRB-IB. In accordance with the French Public Health Code (articles L.1243-4 and R. 1243-61), the CRB-IB has received accreditation from the French authorities to use samples for scientific research. Samples used in the ICGC cohort were collected prospectively by the French Sarcoma Group as part of the ICGC program (International Cancer Genome Consortium). Clinicopathological data and patient information are summarized in **Table S2**. All cases were systematically reviewed by expert pathologists of the French Sarcoma Group according to the World Health Organization guidelines [22]. Patients’ written informed consent was approved by the Committee for the Protection of Individuals. All samples were collected before treatment.

#### Cell lines and primary culture

All cell lines OC80 (LMS; Male), OC88 (LMS; Male), OC48 (LMS; Female), OC98 (UPS; Male) and OC110 (UPS; Female) were primary cultures established as previously described [23] and were cultured in RPMI-1640 (524000-025, Life Technologies, Carlsbad, CA, USA) supplied with 10% fetal bovine serum (S1810-500, Dutscher, Brumath, France). Cells were kept at 37°C in a humidified chamber containing 5% CO_2_.

### METHOD DETAILS

#### Data acquisition

##### Expression microarray data

The 387 complex genetics sarcomas were analyzed on Human Genome U133 Plus 2.0 array (900466, Affymetrix, Santa Clara, CA, USA), according to the manufacturer’s procedures. For GIST, synovial sarcomas, LPS and 87 complex genetics sarcomas, gene expression analysis was carried out by Agilent Whole Human 44K Genome Oligo Array (G4112A, Agilent Technologies, Santa Clara, CA, USA) according to the manufacturer’s protocol.

##### Copy number data

CGH from Affymetrix cohort (**Table S1**: Array-CGH in 53 cases was performed using the BAC-array as described in Chibon et al., 2010 [16], and with Genome-Wide Human SNP 6.0 arrays (901153, Affymetrix, Santa Clara, CA, USA) in 31 cases according to the manufacturer’s protocol with 500 ng DNA as input.

##### Sequencing data

DNA, total RNA and miRNA were extracted from frozen samples of the ICGC cohort and sequenced using Illumina Technologies (Illumina Inc., San Diego, CA, USA) HiSeq2000 for DNA and RNA samples (paired-end) and HiSeq2500 for miRNA samples (single-end). Extraction, library preparation and sequencing protocols are detailed in **Supplementary methods**.

#### Sequencing data analysis

##### RNA sequencing (RNA-seq)

Alignment and expression quantification were performed as previously described [19]. Fusion transcripts were detected with Defuse v0.6.1 [24] as previously described [25]. *miRNA sequencing (miRNA-seq)*

Reads were trimmed for adaptors using Cutadapt version 1.10 [26] with -q 30 and -m 18 parameters. Sequencing quality was assessed using FastQC from the Babraham Institute (https://www.bioinformatics.babraham.ac.uk/projects/fastqc/). We then aligned the reads with mature miRNA sequences from the miRbase Sequence database [27] according to the recommendations in [28]. We first used the BWA-aln algorithm version 0.7.17 [29] with -n 1 -o 0 -e 0 -k 1 -t 8 parameters and evaluated mapping quality using Qualimap version 2.2.2b [30]. We then used Samtools version 1.9 [31] to discard the reads with a mapping quality under 30 and to count mapped reads with the reference sequences (*samtools idxstats*).

##### Whole Genome Sequencing (WGS)

DNA reads were trimmed of the 5’ and 3’ low quality bases (phred cut-off 20, max trim size 30 nt) and sequencing adapters were removed with Sickle2 [32] (https://github.com/najoshi/sickle) and SeqPrep3 (https://github.com/jstjohn/SeqPrep), respectively. Then, DNA-curated sequences were aligned using Bowtie v2.2.1.0 [33], with the very sensitive option, on the Human Genome version hg19. Thus, aligned reads were filtered out if their alignment score was less than 20 or if they were duplicated PCR reads, with SAMtools v0.1.19 [31] and PicardTools v1.118 [34] (http://broadinstitute.github.io/picard/), respectively.

#### Detection of single nucleotide and structural variants

##### Single Nucleotide Variant (SNV)

SNV were detected in RNA-seq and WGS data using Samtools mpileup (SAMtools v0.1.19 [31]), with a minimum of 20 as phred quality score (-Q 20), and bcftools (SAMtools v0.1.19 [35]) with options view –cvg for RNA-seq data and call –Am for WGS data. RNA-seq detected variants with fewer than 5 coverage reads were filtered out. The variants detected in normal, tumor DNA and tumor RNA were merged in the same file. Then, somatic variants were extracted with: (i) a minimum coverage of 14 reads in the tumor and 8 in the normal and (ii) a minimal allelic fraction of 0.3 in tumor and 0 in normal. Variants were annotated using the Annovar v20160314 tool [36]. Variants were selected whose alternative allele frequency (AF) in the Caucasian population (CEU) is lower than 0.1%, as reported in the 1000Genome database [37]. Finally, variants were kept if they were localized in coding regions and were non-synonymous.

##### Structural variants

Breakpoints were detected from WGS data. Paired-end reads were aligned using Bowtie v2.2.1.0 [33], a very sensitive local option allowing soft-clipped sequences. The algorithm has three main steps: i) identification of potential breakpoints, ii) characterization of the second side of the breakpoints, and iii) selection of high-confidence breakpoints. All parameters were set to analyze 60X tumor and 30X normal sequencing depth. Very conservative filters were used to minimize false positive detection. Details are available in the **Supplementary methods**.

Copy number variants (CNV): Genome-Wide Human SNP 6.0 arrays were analyzed as previously described in [38]. Genes absent in more than a third of the patients were discarded. WGS paired tumor/normal data from ICGC were processed using the cn.MOPS R package [39] with default parameters and a 500-nucleotide window. Regions were intersected with TxDb.Hsapiens.UCSC.hg19.knownGene R package version 3.2.2 [40] gene models. Regions with an estimated copy number of 128 were discarded.

For both datasets, genes overlapping segments with different predicted copy numbers were attributed with the lowest number of copies.

#### Experimental validation

##### Fluorescent in situ hybridization

FISH assay was performed on tissue microarrays using the Histology FISH accessory kit (K579911-2, Dako, Agilent Technologies, Santa Clara, CA, USA) according to the manufacturers’ instructions. Thirty-eight tumors from the ICGC cohort were analyzed. Each case was represented by three spots 4 µm-thick and 1mm in diameter. FISH assay was performed using a commercially available MYOCD FISH probe labeled in spectrum orange and a chromosome 17 control probe labeled in FITC (EG-MYOCD-CHR17-20ORGR, Empire Genomics, Williamsville, NY, USA). *MYOCD* and control probe enumeration was performed with a Nikon Eclipse 90i fluorescent microscope with appropriate filters. Pictures were captured using a Pannoramic 250 Flash II Digital Slide Scanner and analyzed with the Pannoramic Viewer (3DHISTECH Ltd., Budapest, Hungary). A case was considered as interpretable when almost 80% of cells presented a signal for both probes. A loss was defined when only one copy of *MYOCD* was observed in the majority of cells; a normal status was when two copies of *MYOCD* were detected in the majority of cells; a gain or polysomy was when 3 to 5 copies of *MYOCD* or both *MYOCD* and the control probe were detected; and amplification was when the number of *MYOCD* signals was equal to or greater than 6, especially when clustered signals were observed.

##### Verification of alterations

For the ICGC cohort, *ATRX*, *TP53*, *RB1*, *PTEN* and *DMD* sequences for each case obtained by whole genome sequencing were entirely screened using the Integrative Genomics Viewer (IGV version 2.6.3 [41]) to search for alterations possibly missed by the detection algorithms used. All SV were verified on gDNA by PCR and Sanger sequencing. MS/NS mutations not found in either WGseq or RNAseq and all FS were verified at both DNA and RNA levels by PCR and RT-PCR, respectively, followed by Sanger sequencing. For samples with enough material left, fusion transcripts detected by RNA-seq were verified by RT-PCR and Sanger sequencing.

To screen mutations on genomic DNA, PCR primers were designed using the Primer 3 program [42](https://bioinfo.ut.ee/primer3-0.4.0/). All PCR were performed on 50ng of DNA using AmpliTaqGold^®^ DNA polymerase (4311820, Applied Biosystems, Foster City, CA, USA) according to the manufacturer’s instructions. PCR program for validating *TP53* mutations is described in [43]. For other PCR, the PCR program used was a Touch-down 60°C program (TD 60°C) (Hybridization temperatures: 2 cycles at a temperature of 60°C, followed by 2 cycles at 59°C, 2 cycles at 58°C, 3 cycles at 57°C, 3 cycles at 56°C, 4 cycles at 55°C, 4 cycles at 54°C, 5 cycles at 53°C and finally 10 cycles at 52°C).

Total RNA was first reverse-transcribed using random hexamers and the High-Capacity cDNA Reverse Transcription Kit (4368814, Applied Biosystems, Foster City, CA, USA) according to the manufacturer’s instructions. All primers used were designed using the Primer 3 program [42](https://bioinfo.ut.ee/primer3-0.4.0). All PCR were performed as previously described for PCR on genomics using the TD60°C PCR program.

##### Sanger Sequencing

PCR products were purified using an ExoSAP-IT PCR Purification Kit (US78200, GE Healthcare, Piscataway, NJ, USA) and sequencing reactions were performed with the Big Dye Terminator V1.1 Kit (4336805, Applied Biosystems, Foster City, CA, USA) according to the manufacturer’s recommendations. Samples were purified using the Big Dye XTerminator Purification kit (4376486, Applied Biosystems, Foster City, CA, USA) according to the manufacturer’s instructions and sequencing was performed on a 3730xl Genetic Analyzer for cohort 1 or 3130xl Genetic Analyzer for cohort 2 (Applied Biosystems, Foster City, CA, USA). Sequences were then analyzed using the Sequencing analysis V5.3.1 and SeqScape V2.6 software (Life Technologies, Carlsbad, CA, USA). FinchTV software (V1.4.0) was also used (Geospiza, Seattle, WA, USA).

##### Immunohistochemistry

IHC assays were performed on tissue microarrays. IHC for PTEN detection was performed on a BenchMark Ultra instrument (Ventana, Washington D.C, USA). Antigen retrieval was performed using a CC1 protocol for 4 min at 100°C (Ventana, Washington D.C, USA). The anti-PTEN antibody (1:200, 9559, RRID:AB_390810, Clone 138G6, Cell Signaling Technology, Danvers, MA, USA) was diluted in Prep kit 26 (783-2876, Roche, Basel, Switzerland) and incubated for 1h. Antibody detection was performed with the Optiview detection kit for 12 min (860-099, Ventana, Washington D.C, USA). IHC for P53 detection was performed on a Bond-III (Leica Microsystems, Wetzlar, Germany) using the clone DO-7 monoclonal antibody (GA61661-2, Dako Omnis, ready-to-use, incubation 20 min, Agilent Technologies, Santa Clara, CA, USA). Antibody detection was performed using EnVision FLEX/HRP (GV80011-2, Dako, Agilent Technologies, Santa Clara, CA, USA). IHC pictures were taken with a Pannoramic 250 Flash II Digital Slide Scanner and analyzed with the Pannoramic Viewer (3DHISTECH Ltd., Budapest, Hungary).

##### Immunofluorescence

Immunofluorescence was performed on tissue microarrays. First, tissues were de-paraffinized in three xylene baths for 5 minutes and then rehydrated in successive baths of ethanol from 100% to 70%. For heat-induced epitope retrieval, slides were incubated for 20 minutes in a microwave oven in DAKO Target Retrieval pH6 (S203130-2, DAKO, Agilent Technologies, Santa Clara, CA, USA). Then, they were incubated with primary antibodies overnight at 4°C in a humidity chamber and after with secondary antibodies for 1 hour at room temperature. Slides were then mounted with Vectashield mounting medium plus DAPI (H-1200-10, Vector Laboratories, Burlingame, CA, USA). Images were acquired on a Zeiss Cell Observer Microscope (Zeiss, Oberkochen, Germany) or a confocal microscope LSM 780 (Zeiss, Oberkochen, Germany). Primary antibody used for Dp427 was the anti-human Dystrophin form Leica Biosystems (1:25, Dy4/6D3, NCL-DYS1, RRID:AB_442080, Leica Biosystems, Wetzlar, Germany) and the Dystrophin antibody from Abcam (1:100, ab15277, RRID:AB_301813, Abcam, Cambridge, UK) was used for all the dystrophin isoforms. Secondary antibodies used were goat anti-mouse Alexa Fluor 488 (1:400, A-11001, RRID:AB_2534069, Thermo Fisher Scientific, Waltham, MA, USA) and goat anti-rabbit Alexa Fluor 594 (1:400, A-11072, RRID:AB_2534116, Thermo Fisher Scientific, Waltham, MA, USA).

##### Cytotoxicity analysis

LMS and UPS cells were seeded in 96-well microplates at a density of 2.10^3^ cells per well in 100µL RPMI-1640 medium (524000-025, Life Technologies, Carlsbad, CA, USA). After 24h incubation at 37°C, 100µL of medium were added with 2X of final concentration (100 to 1.5 µg/mL) of either CCG-1423 (10010350, CAS:285986-88-1, Bertin Bioreagent, Montigny le Bretonneux, France) or CCG-100602 (10787, CAS:1207113-88-9, Bertin Bioreagent, Montigny le Bretonneux, France). Cells were incubated at 37°C for 72h, after which 20µl (5mg/mL) solution of MTT (M2128, 5mg/mL, Sigma, St Louis, MO) dissolved in water were added. After 2h of incubation, media were removed and the MTT metabolic product formazan was dissolved in 100µl DMSO (5879, Sigma, St Louis, MO, USA). Absorbance at 570nm and 650nm was measured with the Clariostar plate reader (BMG Labtech, Ortenberg, Germany) and cell viability was analyzed as follows:

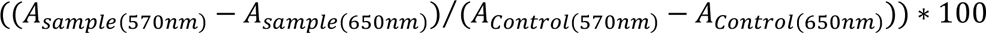

IC_50_ was calculated with GraphPad Prism (GraphPad Software, San Diego, CA, USA) using non-linear regression. Each experiment was done three times in triplicate.

#### Quantification and statistical analysis

##### Normalization of Affymetrix and Agilent micro-arrays and gene selection

We used 87 samples analyzed on Agilent and Affymetrix platforms (**Table S1**). We selected the genes with Pearson’s correlation coefficient (PCC) with itself between chips over 0.8 or better than with any other genes in both experiments. We then normalized gene expression, first in separate experiments and then on a merged dataset by applying quartile normalization (preprocessCore R package version 1.48.0 [44]). We harmonized the expression between the platforms by gene expression median centering in each experiment and then adding the mean of the experiment medians. The method is illustrated in Figure S1 and detailed in Supplementary methods.

##### Gene module clustering

To define groups of co-expressed genes, we computed the pairwise Pearson’s correlation coefficient (PCC) of gene expression with variance > 2 across patients. We built a graph of co-expression with correlated genes (PCC > 0.7) and searched for communities using edge.betweenness.community from igraph R package version 1.2.5 [45].

##### Sample clustering and PCA analysis

In all cases concerning unsupervised clustering, samples were clustered using the *PCA* and *HCPC* functions from the FactoMiner R package version 2.3 [46] and were visualized using the pheatmap R package version 1.0.12 [47]. PCA analysis on ICGC and TCGA miRNA transcriptome was computed using the *prcomp* R function and visualized using the ggbiplot R package [48]. We used the R package Rtsne version 0.15 [49] to visualize GTEX data with the parameters dims=2 perplexity=100 and max_iter=1000.

##### Patient classification

We computed centroids with citccmst R package version 1.0.2 [50] for hLMS and oLMS using the 1672 differentially expressed genes detected in the Affymetrix cohort. Distance to centroids was computed as 1 - Spearman’s correlation coefficient for each patient from all three cohorts. Patient classification was performed using the mclust R package version 5.4.6 [51]. We selected the Gaussian mixture distribution estimation that best fitted the hLMS centroid distance distribution (maximization of Bayesian Information Criterion).

##### Clinical enrichment

Clinical enrichment significance was established using the two-tailed Fisher’s exact test for categorical data comparing one category to the others and Wilcoxon’s test for continuous data.

##### Survival analysis

Survival analysis was performed using the survival R package version 3.1-12 [52] by fitting a simple Kaplan Meier model in which significance was set at log-rank test p-value < 0.01. Survival curves were plotted with survminer version 0.4.6 [53].

##### Differential expression analysis

*mRNA:* Differential expression (DE) analysis was performed using the two-tailed Welch Student’s test corrected for multi-testing using Holm’s method, and t-scores were stored as a measure of hLMS/oLMS expression. Expression of the *MYOCD* gene was considered high if its expression was over the third quartile of the global expression distribution (from all genes in all samples) separately in each experiment.

##### ICGC miRNA

We applied the edgeR R package version 3.28.1 [54] on raw counts to normalize data and define significantly differentially expressed genes with a generalized linear model fitting (*glmQLFit*), correcting the returned p-values using Holm’s method. We kept as expressed any miRNAs (*filterByExpr*) with a summed raw count over all samples > 10 and represented by a minimum of five reads in at least one sample.

##### TCGA miRNA

We computed the differential expression on TCGA miRNAs using the miRComb R package with the *limma* method [55]. We kept miRNAs with a median normalized count >1 in at least one of the LMS groups in both cohorts, having an hLMS/oLMS absolute logFC > 1, and a Holm’s adjusted *limma* p-value < 0.01. We used the *lm* function from the stats R package to compute the R^2^ value between ICGC and TCGA logFC.

##### Functional enrichment and mapping

Modules of co-expressed genes were analyzed using the *enricher* function from the ClusterProfiler R package version 3.14.3 [56] with the Molecular Signatures Database version v6 (MSigDB)[57]. Significance threshold of the hypergeometric test FDR adjusted p-value was set at < 0.05.

Differentially expressed genes were analyzed using the command line version of the GSEA software (version 3.0)[57] and MSigDB v6. We submitted the gene list ranked by hLMS/oLMS t-scores to the *xtools.gsea.GseaPreranked* function with default parameters. Significance threshold on permutation test FDR-adjusted p-value was set at 0.05. For the sake of clarity, only terms with adjusted p-values < 0.01 are reported. Enrichments for regulatory elements in groups of over- and under- expressed genes were performed on the iCistarget webserver (https://gbiomed.kuleuven.be/apps/lcb/i-cisTarget/)[58]. Significance threshold of normalized enrichment score (NES) was set at > 3 by default. Annotations related to position weight matrix predictions from the same transcription factors were grouped together.

We used the GSAn webserver (https://gsan.labri.fr/)[59] with default parameters to exhaustively annotate genes with the most precise Gene Ontology term.

##### miRNA-mRNA interaction analysis

The MiRComb R package [55] was used to integrate miRNA, mRNA expression data and experimentally validated miRNA-mRNA interactions from miRecords v4 [60] and miRTarBase v7.0 [61]. We retrieved 6262 known interactions which represent interactions between 30 DE pre-miRNAs and 3850 genes. Pre-miRNA expression was estimated by averaging signals from derived mature miRNA. We kept interactions for which mRNA and miRNA had an hLMS/oLMS absolute log Fold Change (logFC) > 1, a *limma* p-value < 0.01, a significant Pearson’s anti-correlation (adjusted p-value < 0.01), and were described in at least one of the two databases.

##### CNV recurrence analysis

Alteration recurrence was estimated by computing the frequency of each event (homozygous, heterozygous deletion, gain of one copy and amplification, *i.e.* gain of four copies or more), *i.e.* the number of a given event divided by the total number of patients (missing data being discarded). To evaluate alteration enrichment in each LMS group, losses were grouped into homo- and heterozygous deletions and gains and amplifications (one-tailed Fisher’s exact test p-value < 0.01). We computed enrichment for each type of event in the 291 cytobands by comparing the number of significantly altered genes defined above for each band (one-tailed Fisher’s exact test corrected with Holm’s method < 0.01). If more than one type of event was enriched, the most significant was kept.

##### Mutational pattern analysis

We analyzed patterns of somatic mutation using the MutationalPatterns R package version 1.12.0 [62]. We first generated a 96 tri-nucleotide mutation count matrix per patient which we compared to the 30 COSMIC signatures v3.1 [63]. We kept signatures with cosine similarity > 0.75 and computed the optimal contribution that best explained the observed mutational profiles in patients.

Tumor mutation burden was computed using the total number of somatic variants divided by the total length of human genome version hg19 (22 autosomal and 2 sexual chromosomes).

## Results

### Identification of a group of 42 LMS behaving as simple genetic sarcomas

To detect LMS molecular subtypes within sarcoma samples, we combined micro-array datasets obtained on Affymetrix (387 complex genetic sarcomas including 98 LMS) and Agilent platforms (60 GIST, 58 synovial sarcomas, 50 LPS and 87 complex genetic sarcomas) (**Figure S1A**, total = 555 samples). We selected 9066 genes (out of 17854 genes common to both platforms, **Figure S1B**) showing enough consistency to enable merging and normalization of all datasets (see methods and **Figure S1C**).

We assumed that selecting modules of co-expressed genes that potentially group genes with similar functions would lead to more meaningful patient clustering. We detected 15 co-expression modules (out of 54) carrying at least 5 genes from 455 highly correlated genes. Thirteen modules were significantly associated with biological functions and cellular components (*e.g.* immune system activation, cell cycle, skeletal muscle or smooth muscle-related, adipogenesis, extracellular matrix, apical plasma membrane, genomic positional bias, **Table S3A**, **Figure S1D**). We used these 54 modules in a non-supervised approach to cluster the 555 samples and observed a subgroup of LMS clustering together, while the other LMS were mixed with other pleomorphic sarcomas. This LMS subgroup appeared to behave like sarcomas with a recurrent alteration, *i.e.* with a fairly homogeneous transcriptomic program driven by a strong oncogene [15], as observed with GIST, myxoid liposarcomas and synovial sarcomas (**Figure 1A**). We thus hypothesized that this LMS subgroup (41 patients out of the 98 LMS) could be driven by a strong oncogenic program reflected by this specific gene expression profile.

**Figure 1:**
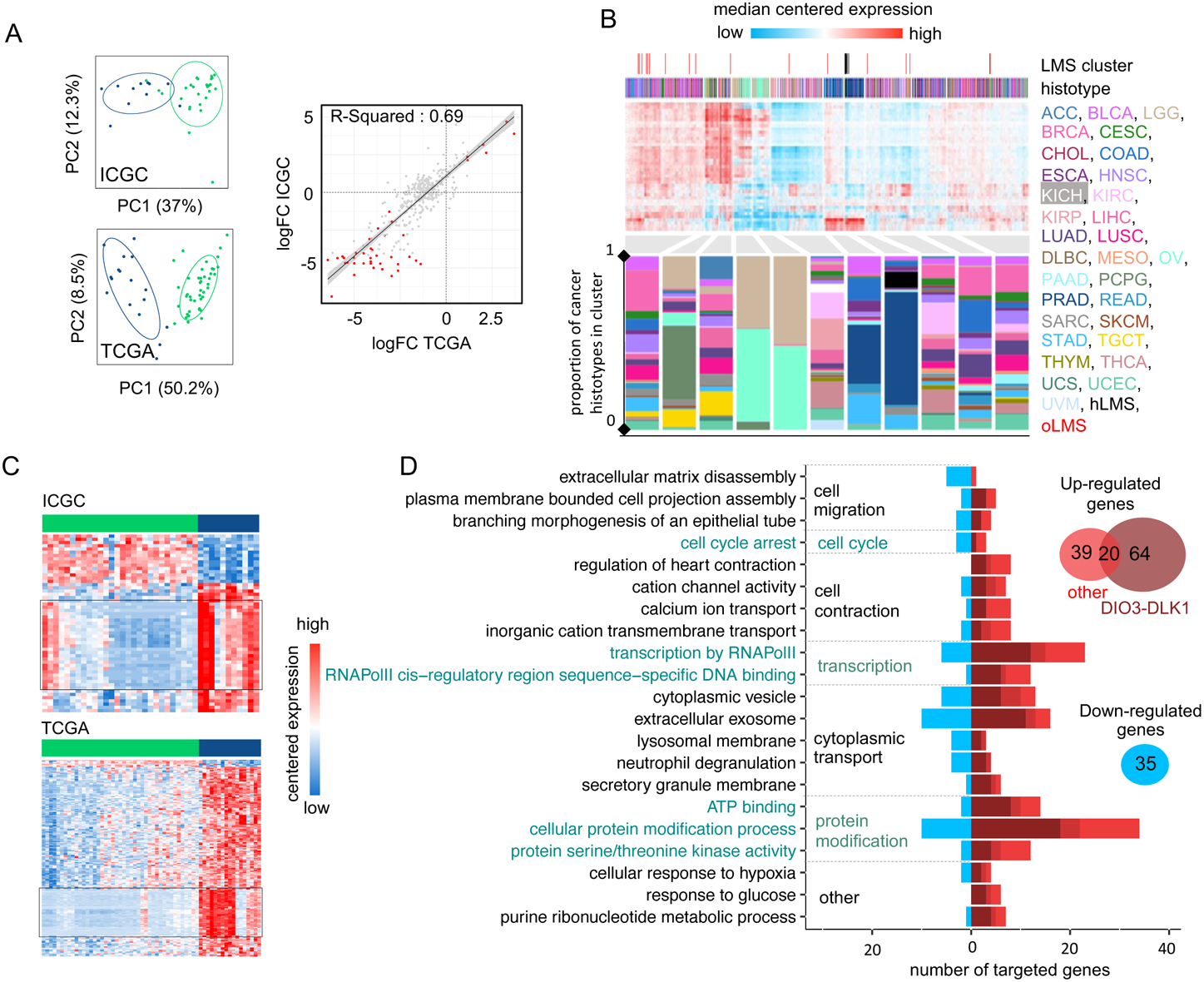
Transcriptional analysis and patient classification: A-B. Heatmaps showing clustering of 555 sarcoma patients and 455 genes (A) and 98 LMS patients and 1672 differentially expressed genes between hLMS and oLMS (B). Patients were clustered using HCPC method and genes are grouped by co-expression modules. Color schemes. Histotype: green forest: leiomyosarcomas, red: GIST, pink: undifferentiated sarcomas, orange: myxoid liposarcomas, blue: dedifferentiated sarcomas, grey: synovial sarcomas, turquoise: other sarcomas. Location: yellow: extremities, green: internal trunk, black: trunk wall. Sex: green: female, blue: male. Grade and differentiation: yellow: 1, green: 2, black: 3. Mitotic count: blue to red: from low to high: A – 0 to 120, B - 0 to 60. Cluster: green: hLMS, blue: oLMS. C. Kaplan-Meier metastasis-free survival analysis in hLMS and oLMS. Number indicates the Log-Rank test p-value. D. GSEA analysis on z-scores obtained from hLMS / oLMS gene expression comparison. Each dot is an enriched term (FDR < 0.01); size corresponds to number of genes involved; x-axis contains mean t-score of all genes annotated in given term and y-axis corresponds to GSEA NES score. Related terms colored the same way. E. i-Cistarget analysis of 843 under- and 800 over-expressed genes in hLMS relative to oLMS. The x-axis represents NES score obtained for over-expressed genes from 0 to the right and for under-expressed genes from 0 to the left. The left and right parts are independent; the enriched features were clustered on the y-axis according to the cell type or tissue they were analyzed from. Histone modifications are only active marks of transcription (H3K4me1, H3K4me3, H3K27ac and H3K9ac). Detailed legends for E-F are available in **Table S3C and S3D** respectively. F. Top panel: distance distribution to centroids (x-axis) computed from transcriptional signature for ICGC and TCGA patients (bars on x-axis). Colors correspond to cluster assignation: patients with a distance lower than 0.6 to one of the centroids were assigned to corresponding centroid (green: hLMS, dark blue: oLMS), while patients with intermediate value were not classified (light grey). Middle and bottom panels: PCA analysis using transcriptional signature genes in ICGC and TCGA cohorts. Each point is a patient, green: hLMS, dark blue: oLMS and light grey: not classified. X-axis and y-axis represent principal components 1 and 2 and their associated representation of variance, respectively.

To select genes that best characterized these LMS, we compared them with the remaining 57 LMS which were mixed with the other sarcomas. As the 98 LMS were all analyzed on the Affymetrix chip, we used the 22635 genes present in the chip. We identified 1672 differentially expressed genes (**Table S4A**) that we used to re-cluster the samples. Almost all samples were classified similarly (95/98) regarding the analysis performed above on the 555 samples. We obtained 42 homogeneous LMS (hLMS) and 56 other LMS (oLMS) (**Figure 1B**).

### hLMS are intra-abdominal, low-grade, metastatic LMS with homogeneous transcriptional behavior

After having confirmed that gene expression profiles within hLMS were significantly more homogeneous than within the other group (Wilcoxon’s test; P = 2.9 x 10^-13^), we tested clinical feature enrichments (**Table 1**). hLMS were mostly located in the abdominal cavity (P = 8.5 x 10^-9^), developed in females (P = 0.003), were well differentiated (P = 3.9 x 10^-9^) and consequently were more frequently grade 1 or 2 (low grades, P = 5.5 x 10^-4^). Interestingly, despite this differentiation and grading, they had a poorer prognosis than oLMS (P = 0.0054, **Figure 1C**).

**Table 1:** Clinical enrichment and gene expression homogeneity in 42 hLMS vs 56 oLMS from Affymetrix cohort and 73 hLMS vs 29 oLMS in combined ICGC (28 vs 11) and TCGA (45 vs 18). (%) indicates that numbers in hLMS and oLMS columns are percentages of patients annotated with first feature (Well: well differentiated, Low: grade 1 + 2, F: female, Internal trunk). M: Male, other: Extremities, Trunk wall, limbs. The p-value was computed using Fisher’s Exact test. Otherwise the median is reported, and the p-value was obtained with Wilcoxon’s test. (+) next to p-values indicates a significant enrichment in hLMS while (-) indicates a significant enrichment in oLMS.

| Feature | Test | Affymetrix |  |  | ICGC + TCGA |  |  |
| --- | --- | --- | --- | --- | --- | --- | --- |
|  |  | hLMS | oLMS | p-value | hLMS | oLMS | p-value |
| Differentiation (%) | Well (vs Poor) | 88 | 24 | $3.87 \times 10^{-9}$ (+) | 84 | 41 | $1.83 \times 10^{-5}$ (+) |
| Grade (%) | Low (vs High) | 58 | 24 | $6.51 \times 10^{-4}$ (+) | | | |
| Sex (%) | F (vs M) | 76 | 48 | 0.003 (+) | 68 | 41 | 0.007 (+) |
| Location (%) | Internal trunk (vs other) | 60 | 7 | $8.48 \times 10^{-9}$ (+) | 82 | 27 | $1.53 \times 10^{-7}$ (+) |
| Mitotic counts (median) | Ranks | 17 | 24.5 | 0.009 (-) | 11 | 35 | 0.0006 (-) |
| Gene expression variance (median) | Ranks | 0.7 | 1 | $2.90 \times 10^{-13}$ (-) | ICGC | | |
| | | | | | 0.25 | 0.36 | $< 2.2 \times 10^{-16}$ (-) |
|  |  |  |  |  | TCGA |  |  |
| | | | | | 0.45 | 0.9 | $< 2.2 \times 10^{-16}$ (-) |

### The transcriptional signature highlights cell cycle and differentiation pathways specific to LMS subgroups

Functional enrichment analysis of differentially expressed (DE) hLMS/oLMS genes (**Figure 1D** and detailed in **Tables S3B** and **C**) revealed biological differences between the two groups. The transcriptional program in hLMS is strongly associated with smooth muscle cell and cell cycle activity, as evidenced by the enrichment of *SRF*, *E2F* and *RB1* targets, CINSARC signature, DNA replication, metabolism and mitochondrial activity in up-regulated genes. In line with these results, activating marks (H3K4me1, H3K27Ac and H3K9Ac) from ChiP-seq experiments in smooth muscles (stomach, rectum, colon, aorta) as well as ChiP-seq peaks for *SRF* and *MEF2A* were enriched in over-expressed hLMS genes (**Table S3D, Figure 1E**). On the other hand, oLMS were associated with unfolded protein response related terms, epithelial-mesenchymal transition and the TGFβ signaling pathway, while enriched histone marks in over-expressed oLMS genes were found to be comparable to those in fibroblasts, epithelial and derived mesenchymal stem cells. These genes are under the regulation of transcription factors (TF) like MYC, ETS1 or ELK1. Therefore, we hypothesize that hLMS and oLMS originate from distinct cell types. To investigate this hypothesis, we stratified patients from two independent multi-omic cohorts.

### Gene signature identifies hLMS in two independent cohorts

To classify LMS from the ICGC (59 patients) [64] and TCGA (75 patients) [13] cohorts, we computed the distance to hLMS and oLMS centroids based on the expression of the 1672 DE genes from the Affymetrix cohort. When the cohorts were merged, 102 cases were strongly enough correlated with one centroid (**Figure 1F**), classifying 73 as hLMS and 29 as oLMS. Computation of clinical enrichment showed hLMS to be mainly intra-abdominal (P = 1.5 x 10^-7^), well differentiated (P = 1.8 x 10^-5^), carried by women (P = 0.007) and with homogeneous transcriptional profiles (P < 2.2 x 10^-16^) (**Table 1**), consistent with information from the training cohort. However, we observed no difference in metastasis-free survival between the two groups.

### hLMS originate from vascular smooth muscle cells

To investigate the potential origin of hLMS, we analyzed the 100 most expressed hLMS genes in 7414 samples from 30 different normal tissues (TCGA GTEX dataset). Using a t-SNE approach, we observed that these genes allowed normal samples to be grouped mainly according to their tissue of origin (**Figure 2**). Visceral smooth muscle tissues were mixed and separated from blood vessels to which the hLMS were the closest. hLMS and oLMS were well separated, and oLMS showed a wider distribution between lung, adipose and breast tissues. These results support our hypothesis that the two LMS groups have a different origin and suggest that hLMS could originate from vascular smooth muscle cells (vSMC).

**Figure 2:**
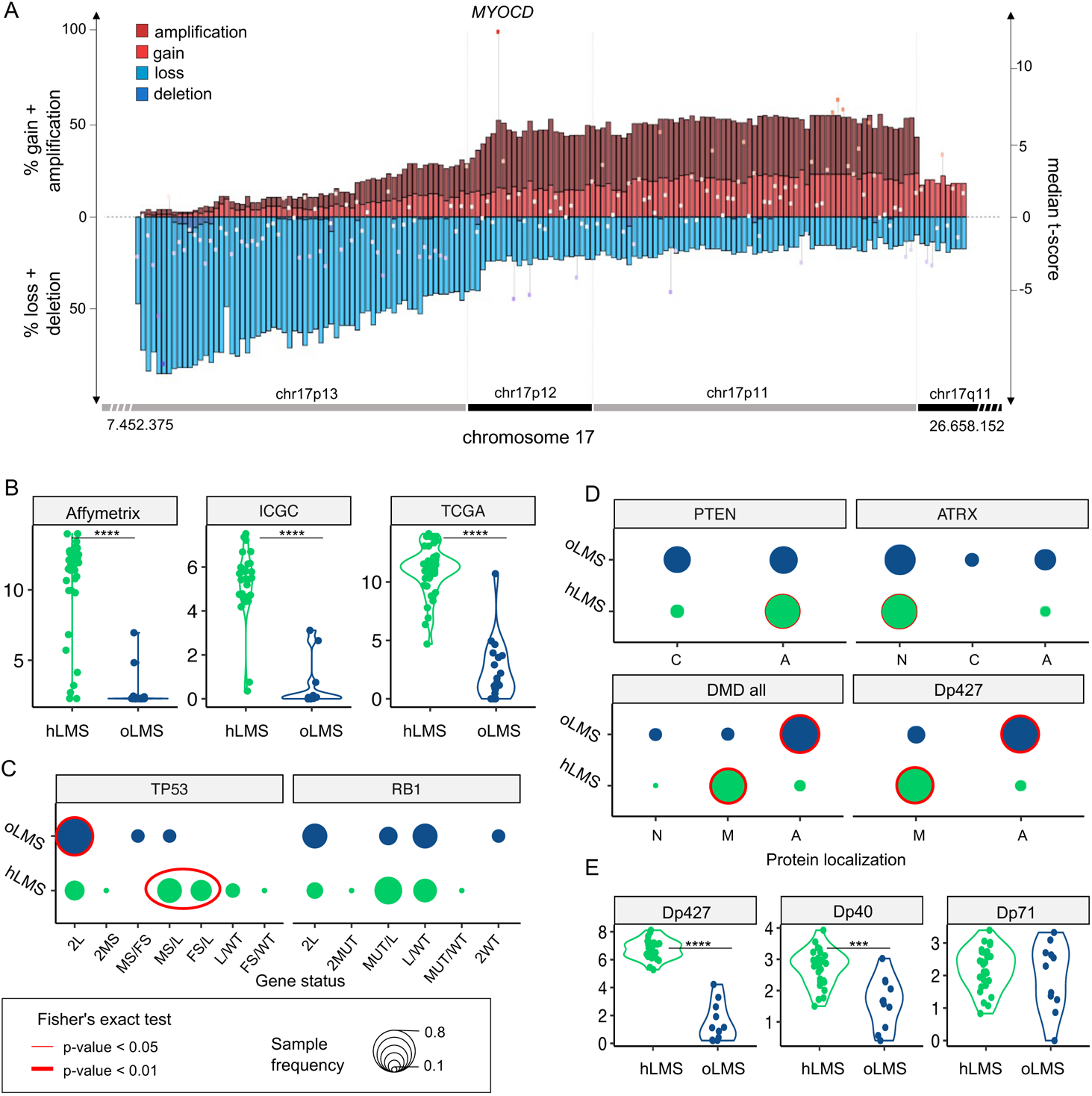
t-SNE clustering from most expressed hLMS 100 genes in 7414 normal and LMS samples from GTEX dataset. Each point represents a sample and the color code corresponds to the tissue type.

We annotated these 100 genes (**Figure S2**) using GSAn [59] (**Table S3E**) and found that 50 of them are part of the extracellular exosome, which are molecules (mRNA or proteins) exported to the extracellular space. This highlights the role of the extracellular matrix (ECM) and of cell-to-cell communication in hLMS pathology. Cell differentiation and migration were represented by 32 and 24 genes, respectively, which suggests the co-existence of both contractile (*MYH11, CNN1, MYL9, LMOD1)* and synthetic (*FN1, TNC, COL1A1/2, MSN, MFAP4*) phenotypes in hLMS.

To complete our analysis of the genomic differences between hLMS and oLMS, we used multi-omics to analyze two additional LMS cohorts.

### miRNAs adopt specific behavior in hLMS

We analyzed 475 expressed mature miRNAs from the 39 patients in the ICGC cohort (28 hLMS and 11 oLMS) and 453 in the 60 TCGA patients (43 hLMS and 17 oLMS). PCA analysis performed with all expressed miRNAs strongly differentiated hLMS and oLMS along the first principal component, which explains 37% (ICGC) and 50.2% of the variance (TCGA) (**Figure 3A**). The high correlation (R^2^ = 0.69, **Figure 3A**) between hLMS/oLMS log-fold changes from both cohorts indicates that each group identified independently in each cohort is consistent and represents two groups of similar diseases. The results of DE analyses of both cohorts are presented in **Table S4B**. We used the TCGA pan cancer (PANCAN) dataset to test our hypothesis that the two groups have a different cellular origin. To this end, we used the 41 significantly differentially expressed miRNAs (35 under-expressed and 6 over-expressed in hLMS) to classify all the cancer samples (**Figure 3B**). All hLMS clustered together among 467 samples mainly from prostate adenocarcinomas (65%), digestive tract tumors (stomach, colon, esophagus, rectum: altogether 14%), LMS (13 gynecological, 8 unclassified: together with hLMS, 13.7%).

**Figure 3:**
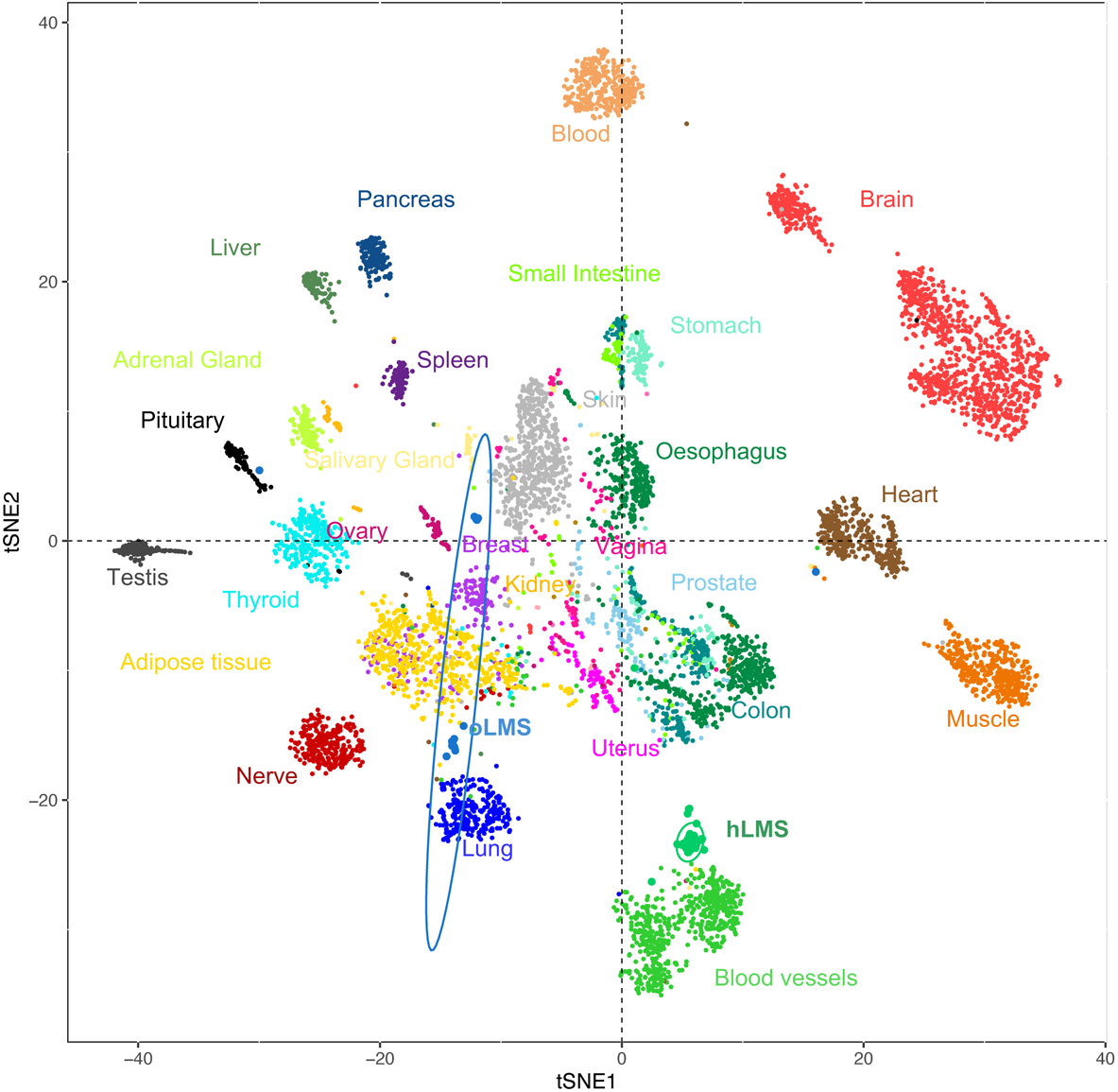
Analysis of miRNA expression **A.** Left panel: PCA obtained from expression of 484 mature miRNAs in 39 ICGC patients (top) and of 475 mature miRNAs in 60 TCGA patients (bottom). Colors correspond to hLMS (green, ICGC: 28 patients, TCGA: 43 patients) and oLMS (blue, ICGC: 11 patients, TCGA: 17 patients). First two principal components are shown with percentage of variance they capture. Right panel: Scatterplot showing correlation between hLMS/oLMS Log Fold Change (LogFC) in ICGC (y-axis) with TCGA (x-axis). Each dot represents a mature miRNA (347 expressed in both cohorts) and red color indicates 71 significant mature DE miRNAs in both cohorts. Line represents linear regression with interval confidence in shaded grey. **B**. HCPC clustering on the 41 mature miRNAs differentially expressed in LMS subtypes across the 9564 PANCAN samples. Heatmap showing median-centered miRNA expression (low: blue to high: red). Column annotations represent histotype of samples (bottom) and focus on LMS clusters (top) for which colors are specified at bottom of figure. Composition in histotype of clusters is detailed in bar plot below heatmap. The y-axis corresponds to the proportion. **C.** Heatmap showing differentially expressed miRNAs (rows) in ICGC (top, 55 miRNAs) and TCGA (bottom, 243 miRNAs). Column annotation corresponds to hLMS (green) and oLMS (blue). Expression values are median-centered (low: blue to high: red). Black rectangles highlight mature miRNAs from DIO3-DLK1 miRNA cluster (ICGC: 26, TCGA: 63, 25 in common). **D.** Functional terms mapped to 158 miRNA targeted genes. The x-axis indicates number of down-regulated (toward left in blue) and up-regulated (toward right in dark red if targeted with only miRNAs from DIO3-DLK1 cluster, medium red if targeted by both miRNAs from DIO3-DLK1 cluster and other miRNAs and light red if targeted by other miRNAs) genes annotated with the term (y-axis).

Interestingly, the eight most discriminative miRNAs of the cluster containing hLMS were 4 over-expressed (*MIR143-3p*, *MIR145-3/5p* and *MIR1*) and 4 under-expressed (*MIR455-3p*, *MIR503-5p* and *MIR424-3/5p*) miRNAs in hLMS that are involved in vascular smooth muscle phenotypic modulation [65–69]. These results corroborated our hypothesis of a smooth muscle origin of hLMS, unlike oLMS which were spread across several clusters.

Intriguingly, all 87 mature miRNAs located in the DLK1-DIO3 imprinted genomic region on chromosome 14 (14q32) were repressed in hLMS. Indeed, 25 miRNAs are among the 35 significantly down-regulated in hLMS (highlighted in **Figure 3C**), 20 others show negative log-fold changes (**Figure S3A**) with very low expression in hLMS (**Figure S3B**), and 42 were not detected in any LMS groups. To evaluate the specificity of this global repression, we compared the expression profiles of the 72 miRNAs (among the 87 DLK1-DIO3) present in the PANCAN dataset. Most hLMS (37/42) clustered within a group of 563 patients, representing 6% of all samples, preferentially with kidney (37%), thyroid (23%), eye (11%) carcinomas and sarcomas (5.5%, 13 gynecological LMS, 1 unclassified LMS, 2 oLMS, 7 UPS, 6 myxofibrosarcomas, 3 dedifferentiated liposarcomas) (**Figure S3C**). hLMS thus cluster with cancer deriving from diverse cell types, which suggests an uncommon repression that might be due to a specific mode of oncogenesis rather than to a vSMC origin.

To evaluate the putative impact of dysregulated miRNAs on hLMS biology, we analyzed their post-transcriptional regulatory network by integrating mRNA and miRNA expression data. We found 210 significant miRNA-mRNA interactions predicted in both ICGC and TCGA cohorts (negative Pearson’s correlation coefficient (PCC), adjusted P<0.01) and present in at least one database (**Table S4C**). We annotated the 158 corresponding target genes (35 down- and 123 up-regulated in hLMS) with GSAn [59]. Twenty-one terms with high specificity were mapped and none of them was specific to the DIO3-DLK1 miRNA cluster, up- or down-regulated target genes, except “response to glucose” which was represented only by up-regulated genes (**Figure 3D,** detailed in **Table S3F**). Dysregulated genes are implicated in major pathways, such as cell migration (“plasma membrane-bound cell projection assembly”, “extracellular matrix disassembly”), cell contraction (“regulation of heart contraction”, “cation channel activity”, “calcium ion transport”), cell cycle and transcriptional regulation.

hLMS miRNA profiles are highly homogeneous and appear to be closely related to vSMC phenotypic modulation, and the predicted regulatory network shows characteristics of both contractile, migratory and proliferative phenotypes. On the other hand, oLMS are heterogenous with no specific program. The question therefore arose whether these two types of LMS also have a distinct genomic mode of oncogenesis.

### hLMS show recurrent and specific genomic instability

LMS are characterized by their highly rearranged genome [17]. Copy number alterations in hLMS appeared more homogeneous than in oLMS both in the merged cohort (**Figures 4A and 4B**) and between the cohorts, which indicates highly correlated penetrance profiles (**Figure S4A**).

**Figure 4.**
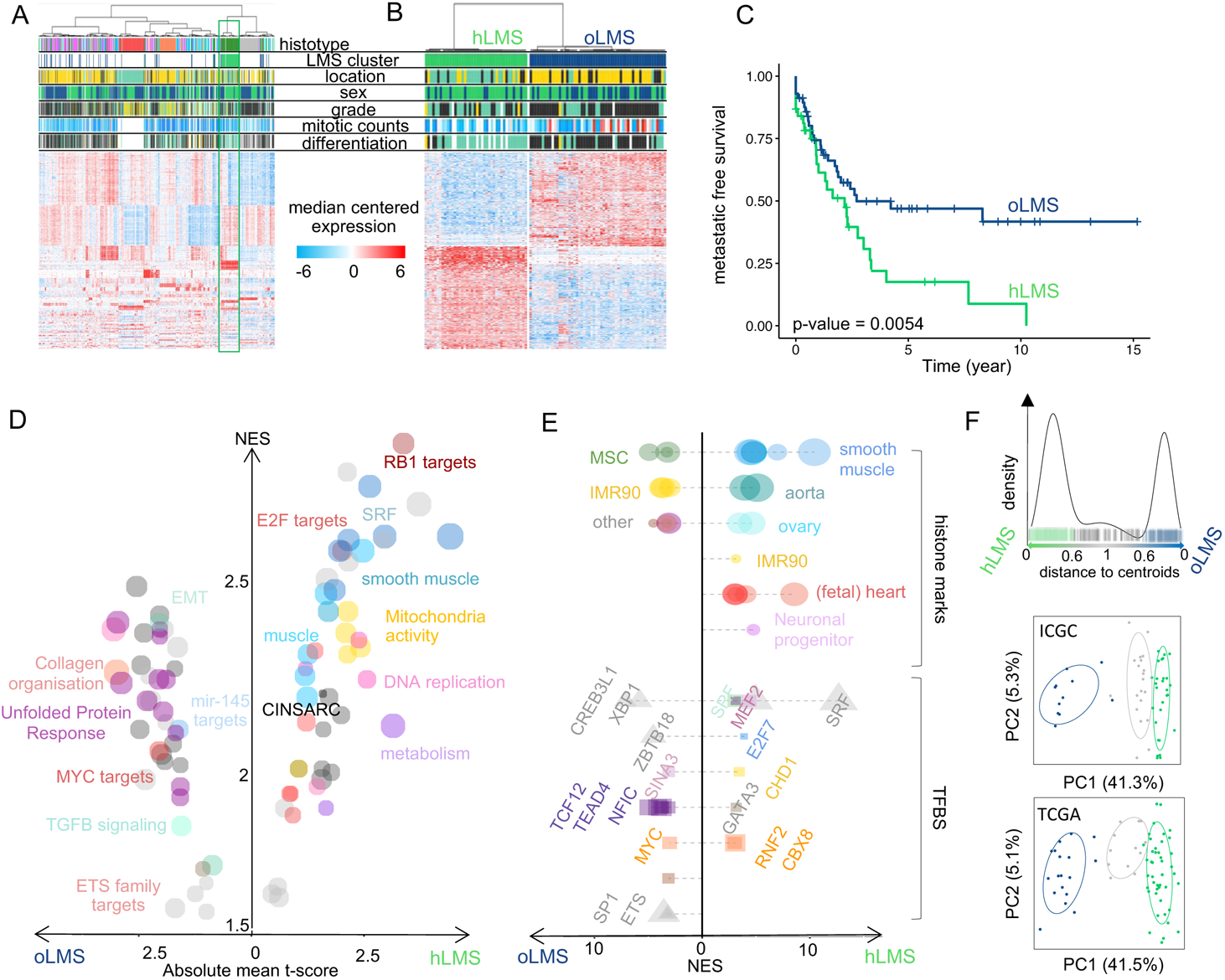
Copy number analysis of 7479 genes in LMS merged cohort (84 Affymetrix (CGH), 39 ICGC (WGS) and 62 TCGA (CGH) patients). **A**. Heatmap showing copy number of genes (columns) in each patient (rows). Patients grouped according to LMS type (hLMS: green, oLMS: blue). Annotations above heatmap show chromosomes from 1 to 22 alternating grey and black and arms (p: green, q: blue). Annotation below shows significantly enriched events in hLMS, Fisher’s Exact test p-value < 0.01. Color scheme is same for copy number and enrichment: homozygous deletion: dark blue, heterozygous deletion: light blue, normal: white, light red: gain of one copy, dark red: gain of 4 or more copies. **B**. Penetrance plot. Percentage (y-axis) of gain (red) and loss (blue) events are represented in hLMS (top panel) and oLMS (bottom panel). Each position on x-axis is a gene that corresponds to genes in A.

Recurrent alterations were significantly enriched in hLMS, especially amplification of chromosome 17p12-p11.2 and loss of chr10q, chr13q14, chr17p13 (**Figure 4B and Table S5A**). The chr17p12-p11.2 amplified region was not only significantly enriched in hLMS, with 31% of hLMS showing an amplification *versus* 7-8% of oLMS but was also the most frequently amplified region in hLMS (**Figure 5A, Table S5A**). Among the genes located in this region, *MYOCD* was the most frequently amplified (36% of hLMS) and was the most over-expressed gene in this region in hLMS compared to oLMS (P < 10^-7^ all cohorts considered, **Figures 5A, 5B, Table S5B**). *MYOCD* expression was very high in 84% of hLMS (97/115, detailed per cohort in **Figure S4B**), whereas it was not expressed or at a very low level in oLMS, even in those with a gain or an amplification (**Figure 5B, Figure S4C**, **Table S6A**). Interestingly, the well-known tumor suppressors *RB1*, *PTEN* and *TP53* belong to three of the eight most significantly enriched lost regions in hLMS: chr13q14 (88% *versus* 72%), chr10q23 (87% *versus* 66%) and chr17p13 (69% *versus* 13%) respectively (all cohorts considered) (**Figures 4, Tables S5A and S5B**).

**Figure 5:**
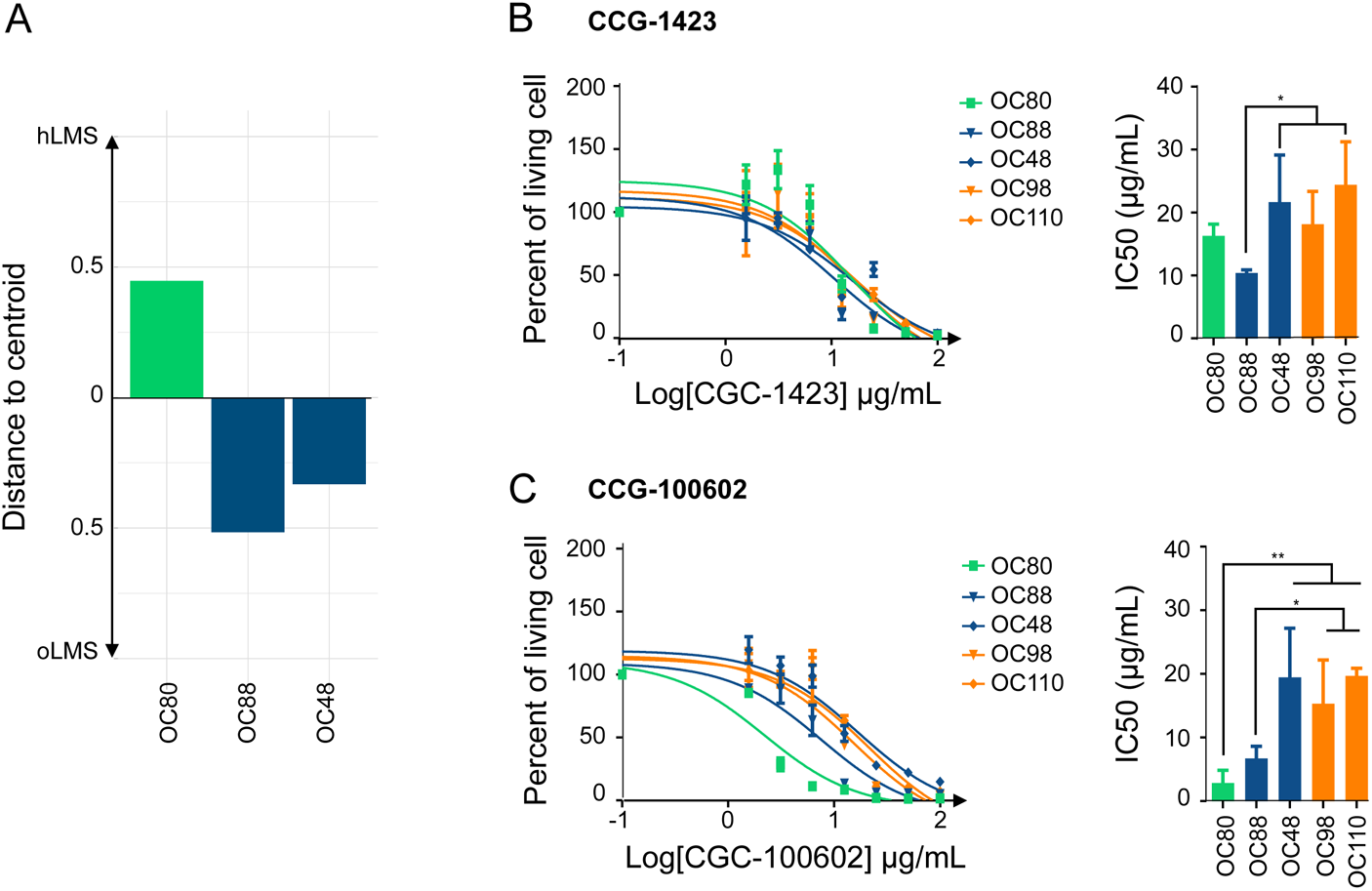
Zoom on genes of interest. **A.** Zoom on chr17p13-11/q11 genomic region (x-axis) penetrance profile containing *MYOCD*. Left y-axis indicates percentage of loss (light blue), deletion (dark blue), gain (light red) and amplification (dark red); right y-axis shows hLMS/oLMS median t-scores (from the three cohorts). Each gene represented by bar (penetrance) and dot (t-score). **B and E**. Violin plots showing expression of *MYOCD* gene and *DMD* isoforms (RNA level, y-axis) in hLMS and oLMS in the three cohorts, respectively. **** indicates a t-test p-value <10^-7^, *** p- value < 10^-3^. **C**. Distribution of *TP53* and *RB1* allele status in hLMS and oLMS. Dot sizes correlate with percentage of patients in LMS group harboring defined status. Cases with biallelic inactivation of *TP53* are compared between groups (2L+2MS: only one mechanism altering both alleles *versus* MS/FS+MS/L+FS/L: two different mechanisms altering each allele): red oval indicates Fisher’s test p-value < 0.01. L: loss, MS: missense, FS: false sense, MUT: MS or FS, WT: wild-type, if 2 is specified, both alleles are concerned. **D**. Cellular distribution of PTEN, ATRX and DMD proteins in hLMS and oLMS. For DMD, localization of its Dp427 isoform is also presented. A: absent, N: nuclear, M: membranous, C: cytoplasmic. Dot sizes correlate with percentage of patients in LMS group harboring defined localization. Red circle indicates Fisher’s Exact test p-value < 0.01 (bold) and < 0.05 (thin).

When we used whole genome characterization of the ICGC to further investigate these genetic variations, we found that oLMS tended to be more rearranged (P < 0.05, **Figure S4D**) than hLMS but that the mutational burden was similar between them (P = 0.5, **Figure S4D**). While no COSMIC mutational signature could be associated with the LMS groups, we found a patient-specific predicted contribution of signatures mainly related to defective DNA repair, except for LMS23, which had a disproportionate mutational burden (120 mutations/Mb *versus* less than 1 mutation/Mb for the other), and a mutational profile similar to ultraviolet light exposure, which is coherent with its location on the scalp. (**Figure S4E**). Very few genes were identified as recurrently mutated (SNV). However, by combining the different alterations, *i.e*. mutations, structural variants (SV) and losses, we found very frequently altered genes across all ICGC-LMS: *TP53* altered in 100% of cases, *RB1* in 97.4%, *PTEN* in 82%, *ATRX* in 28.2% and *DMD* in 25.6%. (**Table 2**, **Tables S6**).

**Table 2:** Summary of genetic alterations in 39 ICGC patients for *TP53*, *RB1*, *PTEN*, *ATRX* and *DMD*. Alterations are categorized as follows: mutation: missense, nonsense, frameshift (FS), non-FS, splicing, SV: structural variant, loss: loss of at least one allele, total: number of patients carrying at least one alteration. Numbers indicate percentage of patients harboring the alteration; the actual numbers are reported between brackets.

| Group | Alterations | *TP53* | *RB1* | *PTEN* | *ATRX* | *DMD* |
| --- | --- | --- | --- | --- | --- | --- |
| hLMS | mutation | 60.7 (17) | 21.4 (6) | 0 (0) | 7.1 (2) | 3.6 (2) |
| oLMS |  | 18.2 (2) | 9 (1) | 0 (0) | 27.3 (3) | 9 (1) |
| all |  | 48.7 (19) | 17.9 (7) | 0 (0) | 12.8 (5) | 5.1 (3) |
| hLMS | SV | 25 (7) | 35.7 (10) | 3.6 (1) | 7.1 (2) | 14.3 (4) |
| oLMS |  | 36.4 (4) | 36.4 (4) | 0 (0) | 0(0) | 36.4 (4) |
| all |  | 28.2 (11) | 35.9 (14) | 2.6 (1) | 5.1 (2) | 20.5 (8) |
| hLMS | loss | 89.3 (25) | 92.9 (26) | 82.1 (23) | 7.1 (2) | 3.6 (1) |
| oLMS |  | 90.9 (10) | 81.8 (9) | 81.8 (9) | 18.1 (2) | 0 (0) |
| all |  | 89.7 (35) | 89.7 (35) | 82 (32) | 10.2 (4) | 2.6 (1) |
| hLMS | total | 100 (28) | 100 (28) | 82.1 (23) | 21.4 (6) | 17.8 (5) |
| oLMS |  | 100 (11) | 90.9 (10) | 81.8 (9) | 45.5 (5) | 45.5 (5) |
| all |  | 100 (39) | 97.4 (38) | 82 (32) | 28.2 (11) | 25.6 (10) |

We found no significant difference in *RB1* and *TP53* global alteration frequencies between the two LMS groups. However, *TP53* presented significantly different alteration patterns. oLMS preferentially lost *TP53* completely (9/11, 82%) whereas 64.3% of hLMS (18/28) exhibited different alterations on each allele with losses, missense and frameshift mutations (Fisher’s exact test, P = 0.01, **Figure 5C, Table S6A**). The same trend was observed for *RB1,* without reaching significance (**Figure 5C, Table S6A**).

*PTEN* was almost exclusively altered by complete gene deletion, regardless of the LMS type (**Table 2**, **Table S6A**). However, although 82% of cases in both groups were altered, its protein expression loss was significantly associated with hLMS (**Figure 5D**).

*ATRX* mutations are described in detail in [64], in which we reported their characterization in the whole ICGC cohort (including the 39 LMS studied here). We showed that *ATRX* alteration and ATRX protein expression loss are associated with uterine LMS and the oLMS type [64]. Accordingly, ATRX nuclear localization was significantly enriched in hLMS (**Table S6A, Figure 5D**).

*DMD* tended to be more frequently altered in oLMS than in hLMS (45.4% and 17.8%, respectively) (**Table 2**). Most *DMD* alterations involved SV, which mainly affects the *DMD* long isoforms (**Tables S6A and S6C**). Regardless of *DMD* genomic status, *Dp427m*, its muscle-specific transcript isoform, was significantly less expressed in oLMS (P = 1.2 x 10^-9^), as was *Dp40* (P = 1.9 x 10^-4^). On the other hand, the expression of *Dp71*, a ubiquitous isoform, was similar in both LMS types (P = 0.63) **(Table S6A, Figure 5E**. Results were confirmed at the protein level, with a significant association of global DMD expression loss and particularly of Dp427 in the oLMS type **(Figure 5D**).

Therefore, despite having similar alteration frequencies of the two major suppressor genes *TP53* and *RB1*, the mechanistic differences and specific expression enrichments of the two LMS types suggest that their oncogenic processes are different. The main features underpinning this distinction are the amplification and strong expression of *MYOCD* and the loss of PTEN protein in hLMS. Indeed, these specific features of hLMS are related to *SRF/MYOCD*, the main drivers of smooth muscle cell differentiation, given that PTEN also interacts with SRF [70].

### hLMS can be targeted specifically with an SRF/MYOCD inhibitor

We tested the hypothesis that the SRF/MYOCD axis could be a driver of hLMS oncogenesis by investigating the therapeutic inhibition of this pathway. We thus studied the impact on cell viability of inhibitors specifically targeting the SRF/MYOCD pathway. CCG-1423, an inhibitor of the SRF/MRTF interaction [71], and CCG-100602, an inhibitor of the SRF/MYOCD interaction [72], were tested on 3 LMS (OC80: hLMS with a *MYOCD* amplification, OC48: oLMS with a *MYOCD* gain and OC88: oLMS) and 2 UPS (OC98 and OC110) cell lines (**Figure 6A**).

**Figure 6:**
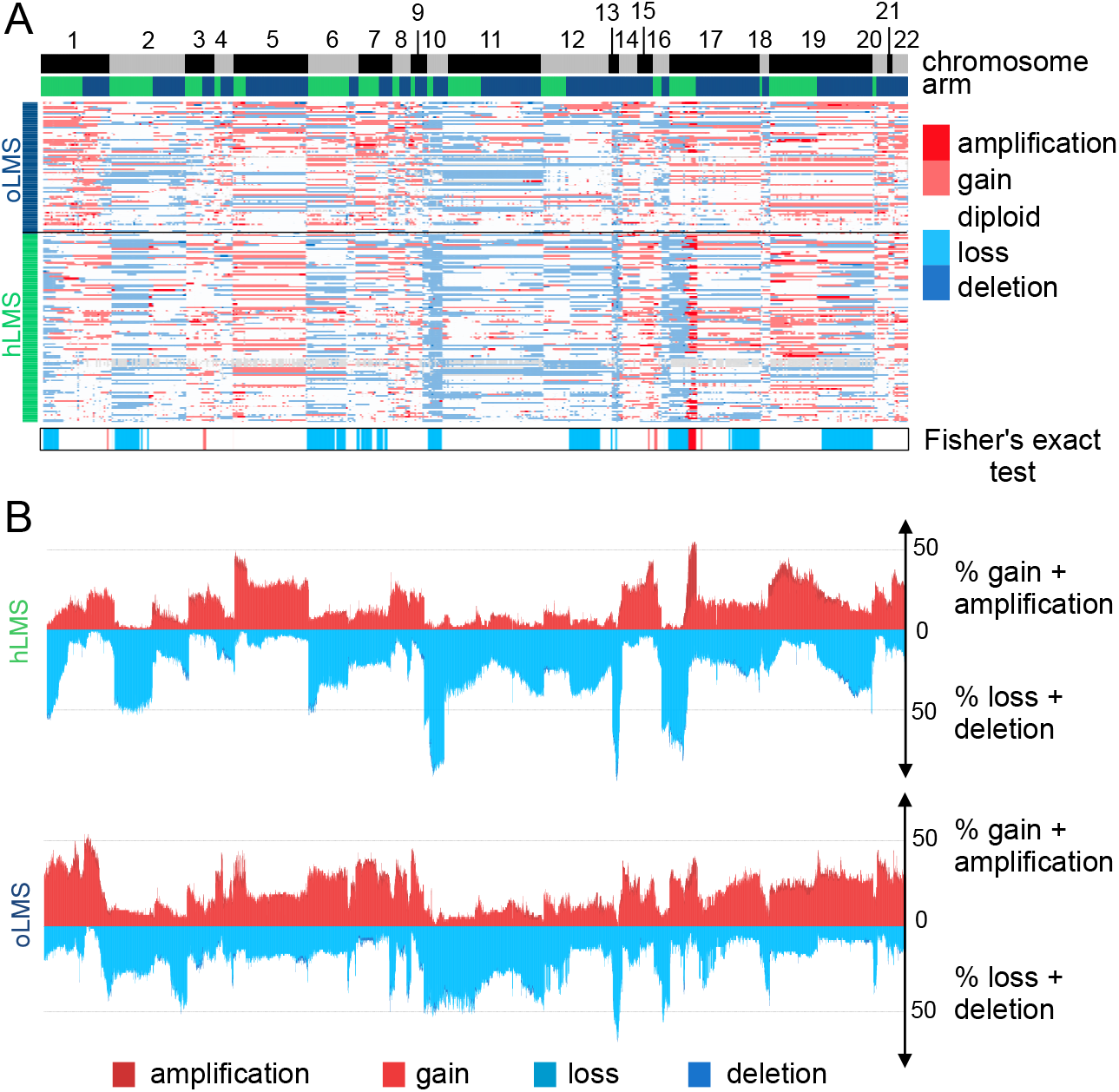
SRF/MYOCD inhibitor can specifically target hLMS. **A.** Distance to centroids determining h/oLMS status on 1672 genes of 3 LMS cell lines. **B**. Cytotoxicity curves of CCG-1423, inhibitor of SRF/MRTF axis, on 3 LMS and 2 UPS cell lines, using MTT assay after 72h of treatment at increasing concentrations (from 1.5 to 100µg/mL). First graph represents one of the three experimentations used to determine IC50 with GraphPad. Second graph shows IC50 (mean ± s.d.; N = 3 independent assays). **C.** Cytotoxicity curves of CCG-100206, inhibitor of SRF/MYOCD axis, on same cell lines, using MTT assay after 72h of treatment at same increasing concentrations. First graph represents one of the three experimentations used to determine IC50 with GraphPad. Second graph shows IC50 (mean ± s.d.; N = 3 independent assays). *p ≤ 0.05, **p ≤ 0.01, ***p ≤ 0.001, p-value was calculated with unpaired t-test for **B** and **C**.

After 72h of treatment with increasing concentrations of each CCG, cell viability assay showed that all cell lines were sensitive to both inhibitors (IC50 ranging from 2.56±1.36 µg/mL to 21.41±3.95 µg/mL) (**Figures 6B** and **6C**). Regardless of their subgroup, LMS were slightly more sensitive to CCG-1423 than UPS, with OC88 reaching significance and being more affected than OC110 (**Figure 6B**). Interestingly, responses to CCG-100602, which is specific to the SRF/MYOCD interaction, exhibited three kinds of behavior: OC80 (hLMS; IC50 = 2.85±1.15 µg/mL) was the most sensitive, OC88 (oLMS; IC50 = 6.70±0.95 µg/mL) had an intermediate response, and OC48 (oLMS; IC50 = 19.44±3.88 µg/mL) and the two UPS (IC50 = 15.32±3.42 and 19.69±0.67 µg/mL) were the least receptive (**Figure 6C**). Overall, all cell lines had a lower IC50 with the SRF/MYOCD inhibitor than with the SRF/MRTF inhibitor. However, the higher responsiveness of hLMS compared to others when SRF/MYOCD was inhibited indicates that the oncogenic dependency on the SRF/MYOCD axis is stronger in hLMS.

## Discussion

Leiomyosarcoma is a very aggressive disease, regardless of its differentiation status, which remains poorly understood. Although previous studies highlighted molecular subgroups and showed the importance of stratifying patients in order to propose targeted and more effective therapies, no consistent biological or oncogenic mechanisms have yet emerged [3–10,17].

By integrating large-scale transcriptomic, epigenetic and genomic data from a large series of LMS, we identified two groups of LMS that are barely similar to the transcriptomic subtypes previously described. In fact, our study differs from the others as we analyzed primary tumors and excluded uterine LMS, unlike most of the other authors [3,6,8,9]. Also, clinical characteristics of each group were similar across the three cohorts, strengthening the signature determining these two LMS sub-groups. hLMS are highly differentiated, preferentially carried by females, low-grade and with an intra-abdominal location, while oLMS are poorly differentiated, high-grade and located in the extremities. Both groups relapsed in at least 50% of cases in all three cohorts and we observed a significant poorer relapse-free survival of hLMS in the Affymetrix cohort. The lack of statistical significance and power in the ICGC and TCGA cohorts may be due to unbalanced group sizes and under-representation of oLMS cases, in contrast with the Affymetrix cohort.

At the molecular level, transcriptomic profiling of oLMS tumors for both mRNA and miRNA highlighted variability within this group, and their genomic alteration patterns were not concordant between the cohorts. Moreover, oLMS showed many regulatory and functional features (active histone marks, many TF binding sites predicted from a cell line showing an epithelial morphology and epithelial-to-mesenchymal related terms) associated with fibroblasts, adipocytes, mesenchymal stem cells (MSC) and epithelial cells, and were spread among normal lung, adipose and breast tissues in the GTEX analysis. Therefore, they could derive i) from the de-differentiation of cells at their location; ii) directly from circulating or local MSC; iii) from fusion between circulating or local MSC with a cell from the tumor site [73]. Yadav et al. [74] showed that cancer cells activate an unfolded protein response (UPR) to adapt to the external (undifferentiated phenotype in a specialized environment) or internal (managing extra material after fusion) micro-environment. Regardless of the origin of oLMS, this mechanism could explain the over-representation of UPR functional terms associated with oLMS over-expressed genes. The heterogeneity observed among patients with oLMS and the difficulty to define a unique oncogenesis might be due to their heterogeneous cellular origin.

In contrast, hLMS seem to have a unique cellular origin closely related to vascular smooth muscles. Indeed, when comparing hLMS with normal tissues, they homogeneously clustered together with blood vessel samples. Moreover, over-expressed genes in hLMS, which are enriched in smooth muscle contractile functions, are mainly associated with active histone marks from smooth muscle datasets. Their promoters were enriched in predicted SRF binding sites (*in silico* and ChIP-seq), which are known to regulate the expression of targeted genes by binding to an element known as the CArG-box, located upstream of smooth muscle (SM) contractile genes [75, 76]. To enhance SM differentiation, SRF needs to cooperate with the MYOCD protein [77], which was found to be over-expressed in more than 84% of hLMS following genetic amplification in 36% of cases. Our data suggest that the over-expression of smooth muscle-related genes in hLMS (compared to oLMS) is triggered by MYOCD, as previously demonstrated by Pérot et al. [10]. Consistently, Dp427, a DMD isoform, which is under the control of MYOCD [78], is nearly always expressed on the membrane in hLMS regardless of *DMD* genomic status, unlike in oLMS in which it is no longer expressed.

Interestingly, the SRF/MYOCD complex regulates all over-expressed miRNAs in hLMS, either directly upon binding to the CArG-boxes present in their promoter [65, 79] or indirectly by targeting their host gene, as in *MIR28* and *LPP,* which has been shown to promote migration in differentiated LMS [10]. More than half of hLMS patients suffer a metastatic relapse. MIR143/145, which are over-expressed in hLMS, are integral components of the SRF/MYOCD network and have been shown to control stress fiber organization and to enhance migratory activity, thereby allowing the cytoskeletal remodeling and phenotypic switching of SMC during vascular injury [69]. MIR143/145 create complex feedback loops by repressing the expression of actin dynamics regulators and SRF/MRTF activity. Xin and colleagues suggested that absence of these miRNAs creates an imbalance and that they are required for SMC cell migration. Additionally, Jiang et al. [65] showed that in differentiated human aortic vSMC, MIR1, which is regulated by SRF/MYOCD and overexpressed in hLMS, suppresses the expression of contractile proteins such as α-SMA and SM22 impairing actin organization, thus creating a negative feedback loop. MIR143/145 and MIR1 may help to fine-tune cytoskeletal homeostasis and allow migration of hLMS tumor cells. hLMS are not only contractile and differentiated (contractile phenotype) but are also proliferative, with migratory features (synthetic phenotype) revealing the co-existence of both phenotypes. These characteristics are those of vSMC, which have highly plastic phenotypes and can cover a wide spectrum of phenotypes from synthetic to contractile [80]. Indeed, markers of both phenotypes, which are highly expressed in blood vessels, are among the most expressed in hLMS. This could represent the natural mixture of vSMC, spanning the phenotypic continuum in these cells. The synthetic phenotype is sustained by the propensity of hLMS to proliferate via strong enhancement of the cell cycle, as suggested by highly frequent *RB1* alterations, the significant over-expression of *E2F1*, the enrichment of up-regulated hLMS genes in E2F/RB1 targets, E2F7 binding sites and cell cycle functional related terms specific to hLMS. This is contradictory with the suggestion of Hemming *et al*. [8], who reported that all LMS possess this feature. It is probably due to the low number of “other” LMS that they had in the different cohorts when comparing LMS with other STS, thus highlighting what they considered to be their “conventional” characteristics.

Although acquired through different preferential mechanisms, alteration of the tumor suppressors *TP53*, *RB1* and *PTEN* occurred equally in hLMS and oLMS and may boost proliferation and cell survival in a non-specific manner. However, the PTEN protein, which is involved in SMC differentiation through direct interaction with SRF [70], is totally absent in hLMS. In fact, the SRF-regulating abilities of SMC are dual: they control the expression of both smooth muscle (SM) contractile genes, depending on MYOCD, and growth-related immediate early genes (IEG) [75], depending on ELK1.

Horita and colleagues showed that in the nucleus, PTEN is directly linked to SRF and MYOCD to form a multi-protein complex and is essential for SRF binding to SM gene CArG-Boxes [70]. Upon forced SMC switch toward proliferation, PTEN is translocated into the cytoplasm, which in turn allows the SRF-cofactor switch and increases IEG gene expression. Therefore, the total absence of PTEN in hLMS may allow SRF to be linked with both SM genes, promoted by the large amount of MYOCD and IEG CArG-Boxes, so SRF may express SMC contractile and proliferative genes. *MYOCD* amplification may overcome PTEN loss and lead to a significantly lower expression of ETS-family target genes in hLMS than in oLMS. Nevertheless, it should be remembered that by comparing two tumor types, the activity of ETS-related transcription in hLMS may be under-estimated. Moreover, the proliferative phenotype of hLMS may be promoted by miRNA repression. Indeed, MIR455, MIR424 and MIR503 under-expression was associated with an increase in pulmonary artery and bladder SMC proliferation by targeting FGF7 [67], CCND1 and CALU [81] and INSR [82] respectively. FGF7 and INSR are thus significantly over-expressed in hLMS and may participate in cell cycle enhancing. Finally, we observed a global repression of the *DLK1-DIO3* miRNAs cluster, which is highly specific to hLMS and papillary thyroid carcinomas. Its predicted target genes are involved in vasculature development and cell migration in both diseases [83] and may play a role in this dual phenotype. However, this cluster involves around 50 miRNAs with different functions in different cellular contexts [84], so understanding how this repression actually impacts hLMS biology requires further investigations.

## Conclusion

Altogether, our findings show that hLMS may originate from vSMC, retain a remarkable degree of plasticity, grow in a maintained differentiated state, and benefit from the enhanced contractile apparatus to migrate. Our data show that the contractile abilities of hLMS come from their vSMC origin rather than arising from an acquired phenotype [10], and suggest that over-expression of *MYOCD* is positively selected, likely triggering tumorigenesis. Accordingly, functional inhibition assays with MYOCD/SRF inhibitors showed efficiency on hLMS, inducing cell death.

## List of abbreviations

LMS: leiomyosacoma
hLMS: homogenous leiomyosarcoma
oLMS: other leiomyosarcoma
GIST: Gastrointestinal Stromal Tumor
PCC: Pearson’s Correlation coefficient
TF: transcription factor
ECM: extracellular matrix
DE: differentially expressed
vSMC: vascular smooth muscle cell
UPR: unfolded protein response

## Acknowledgements

The authors would like to thank the Centre Nacional d’Anàlisi Genòmica (CNAG, Barcelona, Spain) for WG and RNA sequencing services and the Genomics Unit at the Centro de Regulación Genómica (CRG, Barcelona, Spain) for assistance with the smallRNAseq services. The results shown here are partly based upon data generated by the TCGA Research Network: https://www.cancer.gov/tcga. We are grateful to the French Sarcoma Group for tumor banks and associated clinical annotations and to Jean-Baptiste Courrèges. The following French cancer centers also participated in this study: Centre Paul Papin (Angers), Centre Oscar Lambert (Lille), Institut Paoli Calmettes (Marseille). Bioinformatics analyses were performed on the Core Cluster of the Institut Français de Bioinformatique (IFB) (ANR-11-INBS-0013).

## Authors’ Contributions

Conceptualization, Elodie Darbo, Gaëlle Pérot, Lucie Darmusey and Frédéric Chibon; Data curation, Elodie Darbo, Gaëlle Pérot and Lucie Darmusey; Formal analysis, Elodie Darbo, Gaëlle Pérot, Lucie Darmusey, Sophie Le Guellec, Laura Leroy, Laëtitia Gaston, Nelly Desplat, Noémie Thebault, Candice Merle and Frédéric Chibon; Funding acquisition, Jean-Michel Coindre, Jean-Yves Blay and Frédéric Chibon; Investigation, Elodie Darbo, Gaëlle Pérot and Lucie Darmusey; Methodology, Elodie Darbo and Frédéric Chibon; Resources, Philppe Rochaix, Thibaud Valentin, Gwénaël Ferron, Christine Chevreau, Binh Bui, Eberhard Stoeckle, Dominique Ranchere-Vince, Pierre Meeus, Philippe Terrier, Sophie Piperno-Neumann, Françoise Collin, Gonzague De Pinieux, Florence Duffaud, Jean-Michel Coindre, Jean-Yves Blay and Frédéric Chibon; Supervision, Frédéric Chibon; Validation, Elodie Darbo, Gaëlle Pérot and Lucie Darmusey; Visualization, Elodie Darbo, Gaëlle Pérot and Lucie Darmusey; Writing – original draft, Elodie Darbo, Gaëlle Pérot, Lucie Darmusey and Frédéric Chibon; Writing – review & editing, Elodie Darbo, Gaëlle Pérot, Lucie Darmusey and Frédéric Chibon.

## Resource availability

### Lead contact

Further information and requests for resources should be directed to and will be fulfilled by the Lead Contact, Frédéric Chibon

### Materiel Availability

There are restrictions to the availability of human tumor samples due to patient consent. This study did not generate new unique plasmids or reagents.

### Data and Code Availability

Genomic and expression arrays for the other sample will be available on Gene Expression Omnibus (GEO) under accession <u>GSE159847</u> and <u>GSE159848</u> on the 2022-06-30.

ICGC cohort Whole-Genome sequencing and RNA sequencing data for the 67 LMS are available at https://dcc.icgc.org/projects/LMS-FR. miRNA data are available on Gene Expression Omnibus (GEO) under accession <u>GSE159849</u>. Raw files will be available on the 2022-06-30 on Sequence Read Archive under accessions SRP288162. Correspondence between previously published sample identifiers in Gene Expression Omnibus (GSE40021, GSE21050, GSE23980, GSE71118, GSE154591) and in ArrayExpress (E-MTAB-373) datasets and identifiers used in this paper is presented in **Table S1**.

The code is available at https://github.com/ElodieDarbo/lms_onco.

Details for software availability and public datasets are in the supplementary methods.

## Supporting information

### SUPPLEMENTAL FIGURE LEGENDS

**Figure S1 related to methods and Figure 1A:** Normalization of Affymetrix and Agilent micro-arrays.

**Figure S2 related to Figure 2:** Heatmap showing median expression level per tissue type of most expressed hLMS 100 genes.

**Figure S3 related to Figure 3:** Evaluation DIO3-DLK1 miRNA cluster expression.

**Figure S4 related to Figure 5:** Copy number alterations and mutational burden

#### SUPPLEMENTAL EXCEL TABLE LEGENDS

**Table S1 related to Methods:** Presentation of cohort of 555 samples analyzed on micro-arrays and 84 Affymetrix cohort samples for which copy number data were analyzed.

**Table S2 related to Methods:** Clinicopathological characteristics of patients from Affymetrix (98 patients), TCGA (75 patients) and ICGC (59 patients).

**Tables S3 related to Figure 1 and Figure 3**: Results of functional annotations and enrichment in co-expressed modules, differentially expressed genes in Affymetrix cohort, top hundred most expressed genes in hLMS and miRNA target genes.

**Tables S4 related to Figure 3:** Table summarizing differential expression analysis for mRNA in Affymetrix, ICGC and TCGA, for miRNA in ICGC and TCGA and miRComb miRNA-mRNA interaction analysis.

**Tables S5 related to Figure 4:** Table summarizing copy number alteration analysis by cytogenic band and by genes in merged cohort.

**Tables S6 related to Figure 5: A.** Table summarizing genetic alterations occurring for *TP53*, *RB1*, *PTEN*, *DMD*, *ATRX* and *MYOCD* in each sample of ICGC cohort as well as expression data of the six genes. **B.** Table presenting mutations detected in *TP53*, *RB1*, *DMD* and *ATRX*. **C.** Table presenting structural variants detected in *TP53*, *RB1*, *DMD*, *PTEN* and *ATRX*. **D.** Tables presenting primers used on genomic DNA to validate mutations and structural variants. **E.** Table describing primers used on cDNA to validate mutations and fusion transcripts

