## Supplementary Figures for "Distinct cellular origins and differentiation process account for distinct oncogenic and clinical behaviors of leiomyosarcomas"

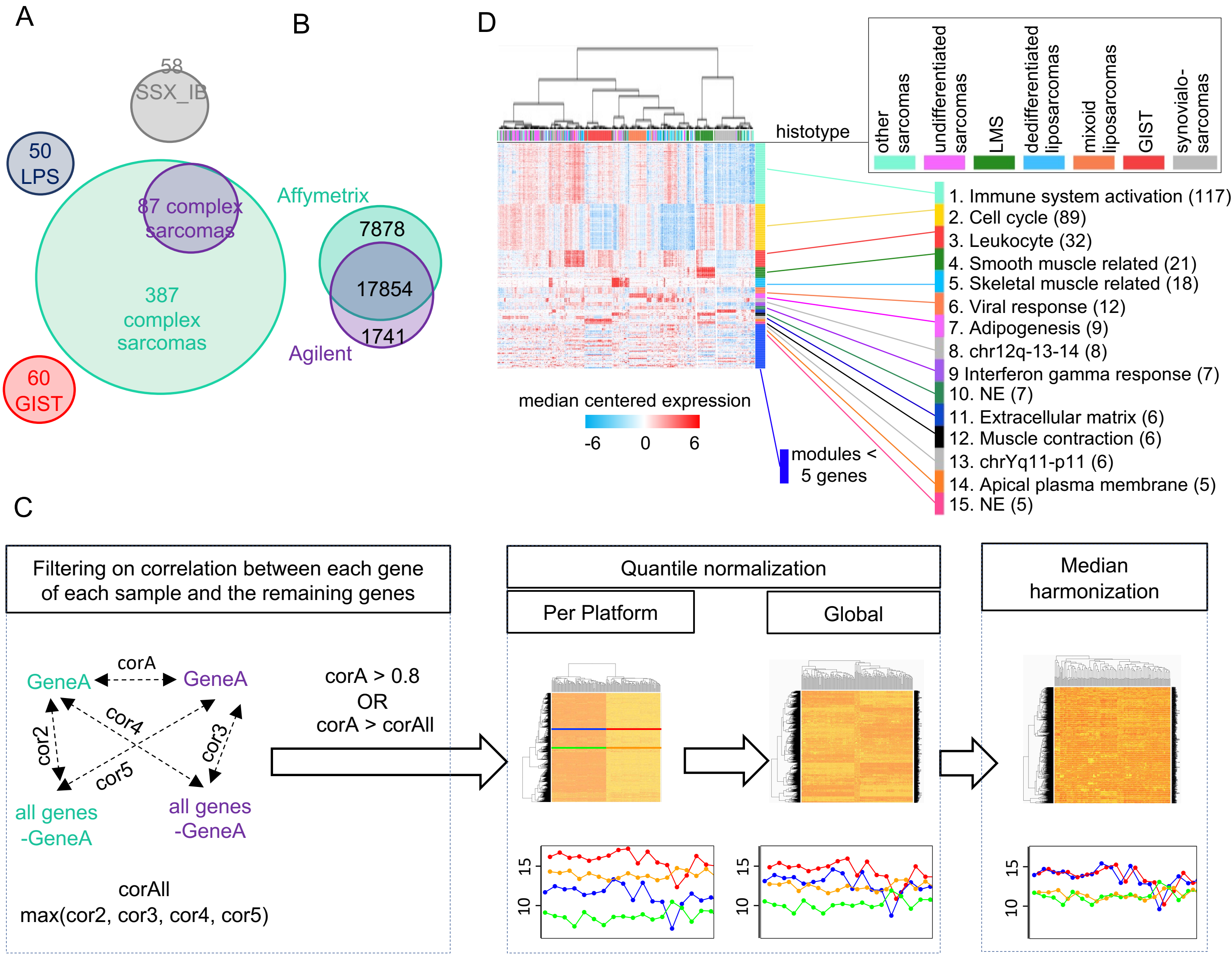


**Figure S1:** Normalization of Affymetrix and Agilent micro-arrays.
**A.** Number of patients per cohort. Overlapping circles represent common samples between experiments. **B.** Venn diagram showing intersection of genes between Affymetrix and Agilent micro-arrays. **C.** Normalization pipeline. The 87 samples analyzed on both Agilent and Affymetrix technologies were used to select the 9066 highly correlating genes between the two platforms (Pearson's correlation > 0.8) or correlating better than with any other genes within and between platforms. The micro-arrays were first quartile-normalized per platform and merged to undergo a second round of quartile normalization. Finally, the median across samples for each gene was computed separately per platform and then harmonization was obtained by subtracting the mean of these medians from the gene expression values. Heatmaps (rows: genes, columns: samples) represent unsupervised clustering of samples at each step of normalization process. The two rounds of quartile normalization cluster the samples per platform, while the harmonized data cluster patients with themselves. Plots under heatmaps show expression profiles (x-axis: expression level) of two randomly chosen genes (gene 1: blue in Affymetrix, red in Agilent, gene 2: green in Affymetrix, orange in Agilent) in the 87 patients. **D.** Summary of functional enrichment analysis of co-expressed gene modules. Heatmap showing clustering of 555 sarcoma patients and 467 genes grouped in modules of co-expressed genes for which a representative functional term is associated. Number before term refers to module number in detailed results (**Table S3A**). Number of genes present in modules is shown in brackets.

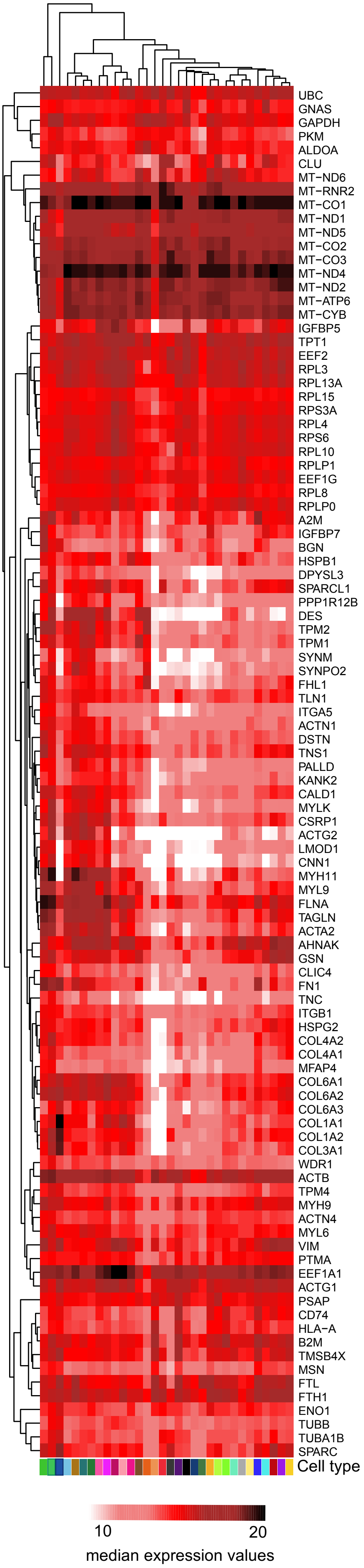


**Figure S2**: Heatmap shows median expression level per tissue type of most expressed hLMS 100 genes.

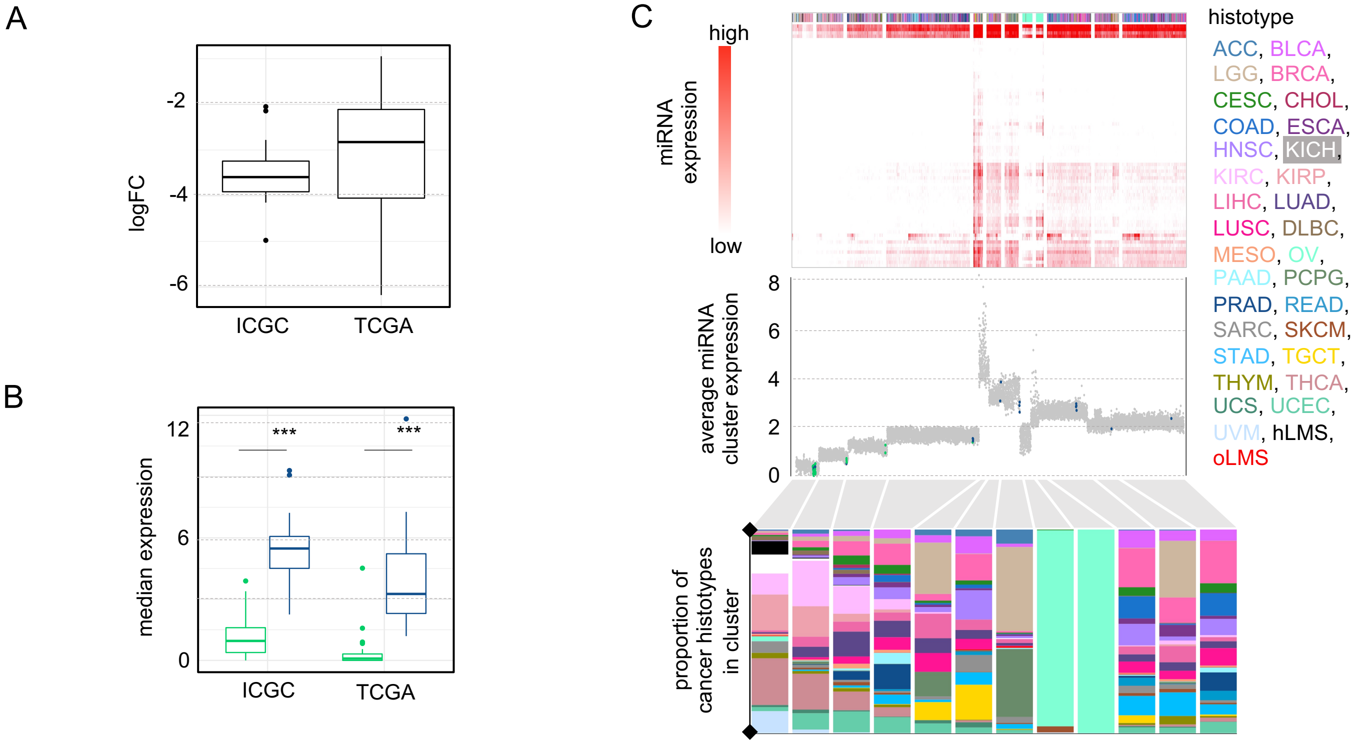


**Figure S3:** Evaluation of 38 DIO3-DLK1 mature miRNAs not DE in either cohort.

**A.** Boxplot representing distribution of LogFC (y-axis). **B.** Boxplot representing distribution of median expression with comparison between hLMS (green) and oLMS (blue). Stars indicate significant difference between LMS groups (Wilcoxon's p-value < 0.0001). **C.** HCPC clustering on 72 mature miRNAs from DLK1-DIO3 miRNA cluster in the 9564 PANCAN samples. Top: Heatmap showing pan-cancer analysis of 83 mature miRNAs from the DIO3-DLK1 cluster in 9564 HCPC-clustered patients. Column annotation indicates histotypes of samples. Middle: Each dot represents a patient ordered on x-axis as in heatmap above. Values on y-axis represent average expression value of the 83 miRNAs in each patient. Green and blue dots indicate hLMS and oLMS respectively.

Bottom: Barplot showing proportion of each histotype in clusters obtained during HCPC analysis.

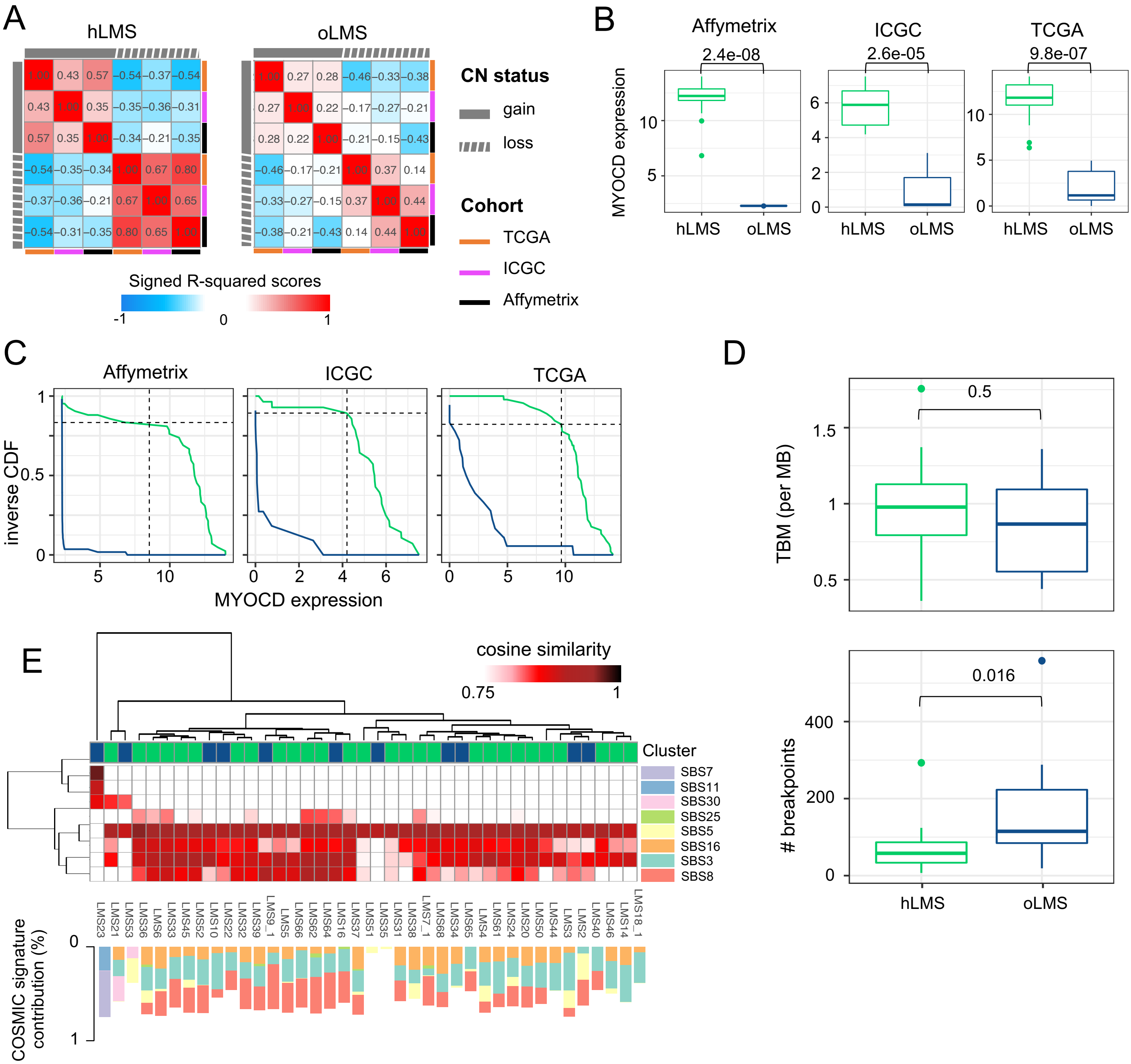


**Figure S4:** Copy number alterations and mutational burden.

**A.** Heatmap showing linear regression r-squared scores representing correlation (anti-correlation: negative, positive otherwise) between gain and loss penetrance profiles of Affymetrix, ICGC and TCGA cohorts. **B.** Boxplot representing *MYOCD* gene expression in patients with *MYOCD* gain or amplification. **C.** Inverse cumulative distribution function (CDF) of hLMS (green) and oLMS (blue) ranked according to *MYOCD* expression in Affymetrix, ICGC and TCGA cohorts. Vertical dashed lines represent 3rd quartile of all gene expressions present in experiments. Horizontal dashed lines indicate proportion of hLMS with *MYOCD* expression which were considered as high expression. **D.** Boxplot distribution of TMB (top: number of somatic mutations per Mb, y-axis) and number of breakpoints (bottom) in hLMS and oLMS. LMS23 was discarded from analysis, TMB=120.35, breakpoints=74. **E.** Heatmap showing cosine similarity between LMS samples (columns, hierarchical clustering: Euclidean distance, average agglomeration) and COSMIC Single Base Substitution (SBS) signatures (rows, hierarchical clustering: cosine similarity, complete agglomeration). Cosine similarity is set to white if under 0.75, and light to dark red (1) otherwise. Columns in bar chart (bottom panel) correspond to same sample as in heatmap. Optimal linear contribution (y-axis) of each SBS signature to reconstruct sample mutational profiles. SBS3: defective homologous recombination DNA damage repair, SBS5: clock-like, SBS7: ultraviolet light exposure, SBS8: unknown, SBS11: temozolomide treatment, SBS16: unknown, SBS25: chemotherapy treatment, SBS30: Defective DNA base excision repair due to *NTHL1* mutation. Numbers above boxplots indicate Wilcoxon's test p-values.
