## Supplementary Methods for "Distinct cellular origins and differentiation process account for distinct oncogenic and clinical behaviors of leiomyosarcomas"

***RNA extraction***

RNA extraction was performed using standard TRIzol (15596026, Thermo Fisher, Waltham, MA, USA) / chloroform extraction (32211-1L, Fisher Scientific, Hampton, NH, USA) followed by 100% ethanol precipitation and RNA purification using the RNeasy Mini Kit (74104, Qiagen, Hilden, Germany) with DNase treatment (79254, RNase-Free DNase Set, Qiagen, Hilden, Germany). Total RNA was quantified using a Nanodrop 1000 spectrophotometer (Thermo Scientific, Waltham, MA, USA) and qualified with an Agilent 2100 Bioanalyzer (Agilent, Santa Clara, CA, USA) using the Agilent RNA 6000 Nano Kit (5067-1511, Agilent, Santa Clara, CA, USA) according to the manufacturer’s instructions.

***RNA sequencing from frozen samples***

To control the sequencing quality, ERCC RNA Spike-In Mix (4456740, Life technologies, Carlsbad, CA, USA) were added to each RNA sample as recommended by the manufacturer. Analysis of this quality control was performed as previously described ^1^.

Libraries from total RNA were prepared using the TruSeq^®^Stranded Total RNA Gold Library Preparation Kit (RS-122-2301, Illumina Inc., San Diego, CA, USA) according to the manufacturer’s protocol. Briefly, 0.5 µg of total RNA was ribo-depleted using the Ribo-Zero Gold Kit. RNA fragmentation resulted in fragments of 80 – 450 nt, with a major peak at 160 nt. First-strand cDNA synthesis by random hexamers and reverse transcriptase was followed by second-strand cDNA synthesis, performed in presence of dUTP instead of dTTP. Blunt-ended double-stranded cDNA was 3´adenylated and Illumina-indexed adapters were ligated. Resulting libraries were enriched with 15 PCR cycles.

Libraries were sequenced on HiSeq2000 (Illumina Inc., San Diego, CA, USA) in paired-end mode with a read length of 2x75 bp using TruSeq SBS Kit v3-HS (FC-401-3001, Illumina Inc., San Diego, CA, USA). Image analysis, base calling and base quality scoring of the run were processed by integrated primary Real Time Analysis (RTA 1.13.48, Illumina Inc., San Diego, CA, USA) software and followed by generation of FASTQ sequence files by CASAVA (v1.8, Illumina Inc., San Diego, CA, USA). Library construction and RNA sequencing were performed at the Centro Nacional de Análisis Genómico (CNAG, Barcelona, Spain).

***miRNA sequencing***

Libraries were prepared by the Genomics Unit at the Centre de Regulació Genòmica (Barcelona, Spain) using the NEBNext Small RNA Library Prep Set for Illumina (E7330L, New England Biolabs Inc., Ipswich, MA, USA) according to the manufacturer’s recommendations. Libraries were sequenced on HiSeq2500 (Illumina Inc., San Diego, CA, USA) in single-read mode with a read length of 50bp (V4 chemistry) to reach a minimal yield of 10M reads.

***DNA extraction***

Genomic DNA from frozen samples of the ICGC cohort was isolated using standard phenol-chloroform extraction protocol. DNA was quantified using Nanodrop 1000 spectrophotometer (Thermo Fisher Scientific, Waltham, MA, USA). Genomic DNA from blood samples was extracted using customized automated purification of DNA from compromised blood samples on the Autopure LS device according to the manufacturer’s recommendations (9001340, Qiagen, Hilden, Germany) with increased centrifugation of 10 min for DNA precipitation and DNA wash.

***Whole genome sequencing***

Whole genome sequencing was performed only on the ICGC cohort. To construct short-insert paired-end libraries, a no-PCR protocol was used with the TruSeq™DNA Sample Preparation Kit v2 (FC-121-2001/FC-121-2002, Illumina Inc., San Diego, CA, USA) and the KAPA Library Preparation Kit (KK8235, Kapa Biosystems, Basel, Switzerland). Briefly, 2 µg of genomic DNA were sheared on a Covaris™ E220, size-selected and concentrated using AMPure XP beads (A63880, Agencourt, Beckman Coulter, Brea, CA, USA) in order to reach a fragment size of 220 – 480 bp. Fragmented DNA was end-repaired, adenylated and ligated to Illumina-specific indexed paired-end adapters.

DNA sequencing was performed in paired-end mode, 2x100 bp or 2x125 bp according to flowcell version, in five or three sequencing lanes of HiSeq2000 flowcell v3 or v4 (Illumina Inc., San Diego, CA, USA) to analyze tumor or normal/constitutive samples and to reach minimal yield of 145 or 85 Gb, respectively. Two tumor samples were sequenced in 20 lanes to reach a minimal yield of 560 Gb. Image analysis, base calling and quality scoring of the run were processed using the manufacturer’s Real Time Analysis software (RTA 1.13.48) and followed by generation of FASTQ sequence files by CASAVA (Illumina Inc., San Diego, CA, USA).

***Breakpoints detection***

The algorithm followed three main steps as detailed bellow.

i) Identification: at this step, reads with at least one soft-clipped end were analyzed as singletons. A position was considered as a potential breakpoint if it was covered by at least 4 soft-clipped reads, 5 soft-clipped bases (with at least two occurrences of two different bases), and if they represented more than 5% of the total amount of reads at this position in the tumor sample. We selected potential somatic events by discarding positions covered by at least 1 read and 1 base in a surrounding 5-nucleotide window in the normal sample. We refer to them as the “first side” of the breakpoint. ii) Characterization: to determine the genomic positions of the soft-clipped sequence from selected reads, we used the UCSC blat server ^2^. If no match was returned, the reverse complement sequence was pulled to test. If there was still no match, the BAM file was investigated for some soft-clip somatic position around the discordant or oversized-insert read mate (hereafter named abnormal) location from the first side of the breakpoint. Because of the small size of the soft-clipped sequence, multiple matches can be found. We used soft-clipped abnormal read mates to select matches with the most coherent chromosomic locations. We refer to them as the “second side” of the breakpoint. iii) Selection: Positions detected from both the first and second sides (in a 5-nucleotide window) were defined as the common pool. We considered as artifacts (due to repeat regions for instance) couples of positions covered with reads and associated soft-clipped sequences separated by fewer than 15 nucleotides and discarded them. We classified the breakpoints in three groups: high-confidence breakpoints, breakpoints needing investigation, and unique position breakpoints. If a breakpoint was covered by reads and associated soft-clipped sequences having both positions belonging to the common pool, it was classified in the first group. If a breakpoint was covered by reads and associated soft-clipped sequences having only one of the positions belonging to the common pool, it was classified in the second group. Then the missing position was searched among the filtered positions. If it was present in the normal sample, the position was discarded and the breakpoint was completed otherwise. Finally, the third group corresponds to breakpoints with both sides outside the common pool and considered as unique: these were discarded. The sides of breakpoints were sorted according to their chromosomic positions to avoid duplicates.

***Normalization of Affymetrix and Agilent micro-arrays and gene selection***

Selecting consistent genes between Affymetrix and Agilent:

${cor}_{max,i}=max\left( cor\left( g_{Affy,i},g_{Agi,\forall j\neq i} \right), cor\left( g_{Affy,i},g_{Affy,\forall j\neq i} \right), cor\left( g_{Agi,i},g_{Agi,\forall j\neq i} \right) \right)$

$$g_{i}=\left\{ 1 Cor\left( g_{i},g_{j=i} \right)>0.8 \bigvee cor\left( g_{i},g_{j=i} \right) > {cor}_{max,i} | 0 otherwise \right\}$$

Where $i,j\mathcal{\in A}\left\{ 1..\mathcal{n} \right\}$ are the indexes of pairs of genes to be compared. $g_{Affy}$ and $g_{Agi}$ are genes analyzed on Affymetrix and Agilent micro-arrays and $\mathcal{A}$ the ensemble of all possible indexes between 1 and n the number of genes to be tested. $g_{i}$ represents the status of gene i that will be kept if 1 or discarded otherwise.

Median harmonization:

$${median}_{i,Affy}=median\left( g_{i,\forall k\in\left\{ Affy \right\}} \right)$$

$${median}_{i,Agi}=median\left( g_{i,\forall k\in\left\{ Agi \right\}} \right)$$

$${median}_{i,All}=mean({median}_{i,Affy},{median}_{i,Agi})$$

$$g_{i,AffyHarm}= g_{i,Affy}- {{median}_{i,Affy}+ median}_{i,All}$$

$$g_{i,AgiHarm}= g_{i,Agi}- {{median}_{i,Agi}+ median}_{i,All}$$

Where $i \in\left\{ 1..\mathcal{n} \right\}$ is the gene index and n the number of genes to be analyzed, $g$ represents the gene expression either in Affymetrix samples ($Affy$) or Agilent samples ($Agi$).

***Visualization***

All generated plots were produced using ggplot2 R package version 3.3.0 ^3^ and extension ggpubr version 0.2.5 ^4^ unless explicitly said otherwise.

**KEY RESOURCES TABLE**

| REAGENT or RESOURCE | SOURCE | IDENTIFIER |
| --- | --- | --- |
| Antibodies | | |
| Rabbit monoclonal anti-PTEN (Clone 138G6) | Cell Signaling Technology | Cat# 9559, RRID:AB_390810 |
| Mouse monoclonal anti-Human Dystrophin (Clone Dy4/6D3 | Leica Biosystems | Cat# NCL-DYS1, RRID:AB_442080 |
| Rabbit polyclonal anti-Dystrophin | Abcam | Cat# ab15277, RRID:AB_301813 |
| Goat anti-Mouse IgG (H+L) Cross-Adsorbed Secondary Antibody, Alexa Fluor 488 | Thermo Fisher Scientific | Cat# A-11001, RRID:AB_2534069 |
| F(ab')2-Goat anti-Rabbit IgG (H+L) Cross-Adsorbed Secondary Antibody, Alexa Fluor 594 | Thermo Fisher Scientific | Cat# A-11072, RRID:AB_2534116 |
| Biological Samples | | |
| GIST (60 samples) | Bergonié Institute | ^5^ |
| Synovial sarcoma (58 samples) | Bergonié Institute | ^6^ |
| Complex sarcomas (278 samples) | Bergonié Institute | ^7^  ^8,9^  ^1^ |
| Complex sarcomas (159 samples) | Bergonié Institute | This paper |
| ICGC cohort of human LMS (frozen sample, FFPE blocs, clinical data) | French Sarcoma Group as part of the ICGC program | This paper |
| Chemicals, Peptides, and Recombinant Proteins | | |
| CCG-1423 | Bertin Bioreagent | Cat# 10010350, CAS:285986-88-1 |
| CCG-100602 | Bertin Bioreagent | Cat# 10787, CAS:1207113-88-9 |
| Critical Commercial Assays | | |
| High Capacity cDNA Reverse Transcription Kit | Applied Biosystems | Cat# 4368814 |
| AmpliTaqGold^®^ DNA polymerase | Applied Biosystems | Cat# 4311820 |
| TruSeq™DNA Sample Preparation Kit v2 | Illumina | Cat# FC-121-2001/FC-121-2002 |
| ERCC RNA Spike-In Mix | Life Technologies | Cat# 4456740 |
| TruSeq^®^Stranded Total RNA Gold Library Preparation Kit | Illumina | Cat# RS-122-2301 |
| NEBNext Small RNA Library Prep Set for Illumina | New England Biolabs Inc. | Cat# E7330L |
| MYOCD FISH probe | Empire Genomics | Cat# EG-MYOCD-CHR17-20ORGR |
| Deposited Data | | |
| Whole-Genome sequencing and RNA sequencing  ICGC cohort | This paper | <https://dcc.icgc.org/projects/LMS-FR> |
| miRNA sequencing data  ICGC cohort | This paper  Availability on 2022/06/30 | Sequence Read Archive: SRP288162  Gene Expression Omnibus: [GSE159849](https://www.ncbi.nlm.nih.gov/geo/query/acc.cgi?acc=GSE159849) |
| Agilent micro-array  Complex sarcomas | This paper  Availability on 2022/06/30 | Gene Expression Omnibus: [GSE159847](https://www.ncbi.nlm.nih.gov/geo/query/acc.cgi?acc=GSE1598497) |
| Agilent micro-array  myxoid LPS | This paper  Availability on 2022/06/30 | Gene Expression Omnibus: [GSE159848](https://www.ncbi.nlm.nih.gov/geo/query/acc.cgi?acc=GSE1598498) |
| Agilent micro-array  GIST | ^5^ | ArrayExpress: [E-MTAB-373](https://www.ebi.ac.uk/arrayexpress/experiments/E-MTAB-373/?query=cancer&page=50&pagesize=250&sortby=raw&sortorder=ascending) |
| Agilent micro-array  Synovial sarcoma | ^6^ | Gene Expression Omnibus: [GSE40021](https://www.ncbi.nlm.nih.gov/geo/query/acc.cgi?acc=GSE40021) |
| Affymetrix micro-array  Complex sarcomas | ^7^  ^8,9^  ^1^ | Gene Expression Omnibus: [GSE21050](https://www.ncbi.nlm.nih.gov/geo/query/acc.cgi?acc=GSE21050), GSE23980, GSE71118 |
| RNA-seq expression (log2+1 RSEM normalised) (TCGA SARC: version 2015-02-24) | ^10^ | RRID:SCR_018938  <http://xena.ucsc.edu/> |
| miRNA expression (TCGA SARC: version 2017-09-08, PANCAN batch corrected: version 2016-12-29) | ^10^ | RRID:SCR_018938  <http://xena.ucsc.edu/> |
| Copy Number Variant (gene level GISTIC2) (TCGA SARC: version 2017-09-08) | ^10^ | RRID:SCR_018938  <http://xena.ucsc.edu/> |
| GTEX+TCGA combined expression data (version 2016-04-12) | ^10^ | RRID:SCR_018938  <http://xena.ucsc.edu/> |
| Clinical annotations (TCGA SARC) | ^11^ | <https://genome-cancer.ucsc.edu/> |
| SNP6 arrays | ^12^ | Gene Expression Omnibus : GSE154591 |
| miRBase (Downloaded Feb 2020) | ^13^ | <http://www.mirbase.org/> |
| miRecords version 4 | ^14^ | RRID:SCR_013021  <http://c1.accurascience.com/miRecords/> |
| miRTarBase version 7.0 | ^15^ | RRID:SCR_017355  <http://mirtarbase.mbc.nctu.edu.tw/> |
| Catalogue Of Somatic Mutations In Cancer (COSMIC) signatures version 3.1 | ^16^ | RRID:SCR_002260  <https://cancer.sanger.ac.uk/cosmic/> |
| Experimental Models: Cell Lines | | |
| OC80 | This paper | NA |
| OC88 | This paper | NA |
| OC48 | This paper | NA |
| OC98 | This paper | NA |
| OC110 | This paper | NA |
| Oligonucleotides | | |
| See supplementary Table S4D and S4E | This paper | NA |
| Software and Algorithms | | |
| Illumina Real Time Analysis | Illumina | RRID:SCR_014332 |
| CASAVA version 1.8 | Illumina | RRID:SCR_001802  <http://support.illumina.com/sequencing/sequencing_software/casava.html> |
| Sickle2 | ^17^ | RRID:SCR_006800  <https://github.com/najoshi/sickle> |
| SeqPrep3 |  | RRID:SCR_013004  <https://github.com/jstjohn/SeqPrep> |
| Bowtie version 2.2.1.0 | ^18^ | RRID:SCR_005476  <http://bowtie-bio.sourceforge.net/index.shtml> |
| SAMtools version 0.1.19, 1.9 | ^19^ | RRID:SCR_002105  <http://htslib.org/> |
| bcfTools version 0.2.0 | ^20^ | https://samtools.github.io/bcftools/ |
| PicardTools version 1.118 | ^21^ | RRID:SCR_006525  http://broadinstitute.github.io/picard/ |
| Defuse version 0.6.1 | ^22^ | RRID:SCR_003279  <https://bitbucket.org/dranew/defuse/src/master/> |
| Cutadapt version 1.10 | ^23^ | RRID:SCR_011841 <https://cutadapt.readthedocs.io/en/stable/> |
| FastQC | Babraham Institue | RRID:SCR_014583 <http://www.bioinformatics.babraham.ac.uk/projects/fastqc/> |
| BWA-aln version 0.7.17 | ^24^ | RRID:SCR_010910  <http://bio-bwa.sourceforge.net/bwa.shtml> |
| Qualimap version 2.2.2b | ^25^ | <http://qualimap.conesalab.org/> |
| Annovar version 20160314 | ^26^ | RRID:SCR_012821  <http://www.openbioinformatics.org/annovar/> |
| Integrative Genomics Viewer (IGV) version 2.6.3 | ^27^ | RRID:SCR_011793  <http://www.broadinstitute.org/igv/> |
| Primer 3 version 0.4.0 | ^28^ | RRID:SCR_003139  <https://bioinfo.ut.ee/primer3-0.4.0/> |
| FinchTV version 1.4.0 | Geospiza | RRID:SCR_005584 <http://www.geospiza.com/Products/finchtv.shtml> |
| Fiji | ^29^ | RRID:SCR_002285  <http://fiji.sc> |
| GraphPad Prism | GraphPad | RRID:SCR_002798  <http://www.graphpad.com/> |
| Gene Set Enrichment Analysis (GSEA) | ^30^ | RRID:SCR_003199  <http://www.broadinstitute.org/gsea/> |
| BLAT | ^2^ | RRID:SCR_011919 [http://genome.ucsc.edu/cgi-bin/hgBlat?command=start](http://genome.ucsc.edu/cgi-bin/hgBlat?command=start" \t "_blank) |
| TxDb.Hsapiens.UCSC.hg19.knownGene version 3.2.2 | ^31^, R package | <http://bioconductor.org/packages/release/data/annotation/html/TxDb.Hsapiens.UCSC.hg19.knownGene.html> |
| edgeR version 3.28.1 | ^32^, R package | RRID:SCR_012802  <http://bioconductor.org/packages/edgeR/> |
| miRComb | ^33^, R package | <https://github.com/mariavica/mircomb> |
| ggplot2 version 3.3.0 | ^3^, R package | RRID:SCR_014601  <https://cran.r-project.org/web/packages/ggplot2/index.html> |
| ggpubr version 0.2.5 | ^4^, R package | <https://CRAN.R-project.org/package=ggpubr> |
| ggbiplot | ^34^, R package | <https://github.com/vqv/ggbiplot> |
| clusterProfiler version 3.14.3 | ^35^, R package | RRID:SCR_016884  <http://bioconductor.org/packages/release/bioc/html/clusterProfiler.html> |
| FactoMineR version 2.3 | ^36^, R package | RRID:SCR_014602  <http://factominer.free.fr/index.html> |
| preprocessCore version 1.48.0 | ^37^, R package | <https://bioconductor.org/packages/release/bioc/html/preprocessCore.html> |
| pheatmap version 1.0.12 | ^38^, R package | RRID:SCR_016418  <https://www.rdocumentation.org/packages/pheatmap/versions/0.2/topics/pheatmap> |
| Rtsne version 0.15 | ^39^, R package | RRID:SCR_016342  <https://cran.r-project.org/web/packages/Rtsne/index.html> |
| citccmst version 1.0.2 | ^40^, R package | <https://CRAN.R-project.org/package=citccmst> |
| cn.MOPS | ^41^, R package | RRID:SCR_013036  <http://bioconductor.org/packages/2.12/bioc/html/cn.mops.html> |
| GSAn | ^42^, online tool | <https://gsan.labri.fr/> |
| iCistarget | ^43^, online tool | <https://gbiomed.kuleuven.be/apps/lcb/i-cisTarget/> |
| igraph version 1.2.5 | ^44^, R package | RRID:SCR_019225  <https://igraph.org/r/> |
| mclust version 5.4.6 | ^45^, R package | <https://CRAN.R-project.org/package=mclust> |
| MutationalPatterns version 1.12.0 | ^46^, R package | <https://bioconductor.org/packages/release/bioc/html/MutationalPatterns.html> |
| survival version 3.1-12 | ^47^, R package | <https://CRAN.R-project.org/package=survival> |
| survminer version 0.4.6 | ^48^, R package | <https://CRAN.R-project.org/package=survminer> |
| R version 3.6 | R Project for Statistical Computing | RRID:SCR_001905  <http://www.r-project.org/> |
| Molecular Signatures Database (MSigDB) version 6 | ^49^, Broad institute | RRID:SCR_016863 [http://software.broadinstitute.org/gsea/msigdb/index.jsp](http://software.broadinstitute.org/gsea/msigdb/index.jsp" \t "_blank) |
| Analytic pipeline | This paper | OCEANCODE: provisional DOI:10.24433/CO.0299110.v1  https://github.com/ElodieDarbo/lms_onco |
